# Digital PCR detection of *Mycobacterium tuberculosis* and HIV-1 co-localization in spinal tuberculosis biopsies

**DOI:** 10.1101/2025.06.11.659223

**Authors:** Robyn Waters, Adrian R Martineau, Maritz Laubscher, Robert N Dunn, Michael Held, Melissa-Rose Abrahams, Anna K Coussens

**Affiliations:** Division of Orthopaedic Surgery, Department of Surgery, Faculty of Health Sciences, University of Cape Town, South Africa; Wellcome Discovery Platforms in Infection, Institute of Infectious Disease and Molecular Medicine, Faculty of Health Sciences, University of Cape Town, South Africa; Centre for Immunobiology, Blizard Institute, Faculty of Medicine and Dentistry, Queen Mary University of London; Division of Medical Virology, Department of Pathology, Institute of Infectious Disease and Molecular Medicine, Faculty of Health Sciences, University of Cape Town, South Africa; Infection and Global Health Division, Walter and Eliza Hall Institute of Medical Research (WEHI), Parkville, Victoria, Australia; Division of Medical Microbiology, Department of Pathology, Faculty of Health Sciences, University of Cape Town, South Africa; Department of Medical Biology (WEHI), Faculty of Medicine, Dentistry and Health Sciences, University of Melbourne, Parkville, Victoria, Australia

**Keywords:** STB, ddPCR, diagnostic, *Mtb*, TB-HIV co-infection, HIV reservoir, biopsy, tissue, bone

## Abstract

**Background:** *Mycobacterium tuberculosis* (*Mtb*) and HIV-1 co-infection in tissues is suggested to favour reciprocal replication, infection and reservoir expansion. Yet, confirmation of this detrimental synergism in diseased tissues is limited.

**Methods:** In this prospective study of 25 adults investigated for spinal tuberculosis (STB) in South Africa (13 (52%) people living with HIV-1 (PLWH), on antiretroviral treatment) 25 open surgery or CT-guided biopsies were collected and portioned into 93 segments. Extracted DNA was analysed by droplet digital PCR (ddPCR) to detect and quantify *Mtb* complex (MTBC) (*rpoB*, *IS6110*), HIV-1 (*pol*, *gag*) and Human (*RPP30*) gene copies. ddPCR sensitivity for *Mtb* was validated against Xpert-Ultra and culture. Total biopsy and intra-biopsy variation in pathogen DNA abundance, co-detection, and relationships to human cellularity, HIV status, and peripheral viral load (VL) evaluated.

**Results:** ddPCR detected MTBC DNA in biopsies from 10/10 (100%) culture-confirmed STB, 5/6 (83%) Xpert Ultra-confirmed STB and 4/9 (44%) diagnosed Not STB (all 4 with previous pulmonary TB). Detected MTBC ranged from 8-59,144 *rpoB* copies/biopsy. *RpoB* copies/million human cells were higher in biopsies from PLWH (p=0.0096) and positively correlated with matched-segment HIV-1 *pol* copies/million cells (r=0.40; p=0.0003), but not VL. HIV-1 DNA was detected in all PLWH biopsies, four with undetectable VL. HIV-1 *pol* copies/million cells were higher in segments with MTBC DNA co-detected (p=0.011) and also correlated with VL (r=0.91; p=0.0003).

**Conclusions:** Reciprocal relationships exists between *Mtb* and HIV-1 abundance in spinal tissue. Findings support investigating TB-HIV co-infected tissue segments to characterise how the immune microenvironment impacts HIV-1/*Mtb* reservoir persistence and/or expansion.

## INTRODUCTION

In persons living with HIV (PLWH), the presence of HIV-1 co-infection can complicate the diagnosis of tuberculosis (TB), with atypical radiographic findings, reduced sensitivity of diagnostic tools^1^, and TB often manifesting in extrapulmonary sites which are difficult to access without invasive procedures.^2^ In South Africa, TB-HIV coinfection rates are high, and development of tuberculosis in the spine (STB) is one of the most common EPTB sites in PLWH.^3–6^ *Mycobacterium tuberculosis* (*Mtb*) infection of the spine, and associated inflammatory damage to the spinal architecture, is painful and debilitating. Rapid diagnosis on biopsy, using highly sensitive techniques, and prompt initiation of anti-tuberculous treatment, are crucial to reduce morbidity and life-long disability in STB patients.^4,7–11^

We and others have previously hypothesized that not only may active HIV-1 infection increase *Mtb* replication at co-infection sites precipitating TB development, but that the inflammatory tissue microenvironment at the site of *Mtb* infection may reciprocally increase HIV-1 reservoir size and activation, by driving viral replication and increasing the pool of infection susceptible cells.^12,13^ Such a relationship may be particularly evident when TB develops within an HIV-1 sanctuary/anatomical reservoir site, given migrating and *Mtb*-specific T cells have increased susceptibility to HIV-1 infection.^14–16^ Counter to the often paucibacillary nature of pulmonary TB (PTB) in PLWH, we recently found PLWH on ART had no significant difference in spinal biopsy total *Mtb* abundance, using surrogate measures of time to culture positivity or Xpert MTB/RIF Ultra cycle threshold (Ct), compared to HIV-1 uninfected patients.^17^ We also found Xpert MTB/RIF Ultra to have a higher sensitivity than culture in detecting *Mtb* in open surgery and CT-guided biopsies.^17^

Digital droplet PCR (ddPCR) utility to detect *Mtb* in clinical samples has recently been demonstrated using DNA derived from blood and formalin-fixed and paraffin-embedded (FFPE) samples from EP sites^18–22^, although utility for the specific detection of *Mtb* in bone and joint biopsies has yet to be explored. Recent findings in CD34+ stem cells from the whole blood of healthy individuals have implicated the bone marrow as a potential niche for *Mtb* infection, with MTBC DNA more frequently detected by ddPCR in circulating CD34+ cells than CD34-, with increased MTBC DNA detection in PLWH.^22^ The bone marrow, as a deep anatomical tissue site, has also been implicated as a viral tissue reservoir that enables the persistence of HIV-1 infection in PLWH on ART^23^, although much remains unclear about the mechanisms of this persistence. HIV reservoir quantification is essential for evaluating HIV cure strategies. Multiplexed ddPCR assays also provide the unique opportunity to quantitate the HIV-1 reservoir, ensuring sensitive, and specific quantification of viral genomes from multiple HIV-1 subtypes including subtype C, which dominates in South Africa and globally, and may thus provide valuable insights into tissue reservoir dynamics following initiation of ART^24–27^ and the impact of and on tissue co-localised *Mtb* infection.

The aim of this study was therefore to optimize and evaluate the utility of ddPCR to detect and accurately quantify *Mtb* and HIV-1 DNA in both open surgery and percutaneous CT-guided needle spinal biopsy tissue, to determine i) if ddPCR is more sensitive than clinically validated tools for *Mtb* detection and quantification in STB biopsies, ii) if HIV-1 proviral DNA can be detected at the site of spinal TB in ART established PLWH, iii) if spinal tissue *Mtb* abundance associates with tissue localised presence and/or abundance of HIV-1, and iv) whether peripheral HIV-1 viral load (VL) is associated with spinal tissue *Mtb* and/or HIV-1 abundance.

## METHODS

### Study design

This is a sub-study of a prospective, longitudinal, observational cohort previously described.^17^ Adults (>18 years) with suspected STB were recruited at Groote Schuur Hospital, a public hospital servicing a population of low socio-economic backgrounds, and low-income households in densely populated vulnerable communities in Cape Town, South Africa, between November 2020 and December 2021. Based on diagnostic results of Xpert MTB/RIF Ultra (GXPU), MGIT TB culture (TBCUL) and histology/histopathology (HIST) from the main study^17^, patients were divided into three groups: *Culture-confirmed STB* (TBCUL+GXPU+HIST+), *Xpert Ultra-confirmed STB* (TBCUL-GXPU+HIST+), *Not STB* (TBCUL-GXPU-HIST-, alternate diagnosis Supplementary Table S1). The study was approved by the Human Research Ethics Committee, University of Cape Town (HREC 606/2019). Written informed consent was obtained from all patients.

### Data and biopsy collection and processing

Clinical data were collected as previously described^17^ and detailed in Supplementary Methods. Spinal biopsy samples were collected via percutaneous CT-guided needle biopsy or open surgery biopsy during mechanical stabilization in theatre, as previously described.^17^ Tissues were immediately submerged in DNA/RNA Shield™ (Zymo Research, diluted 1:1 nuclease-free water (Sigma)), maintained at 4°C overnight, and stored at-80°C. Tissue homogenization workflow, DNA extraction and quantification is detailed in Supplementary Methods and Supplementary Figure S1. Briefly, after thawing, larger open biopsies were cut into four equal sized tissue segments measuring approximately 1 cm^2^ each homogenized independently for DNA extraction. A fifth sample comprising of combined left-over biopsy homogenate from each of the four segments was also prepared for a separate DNA extraction. Small CT-guided biopsy cores, ranging from 2-7 mm long by 1 mm wide, were used in their entirety as a single sample (Supplementary Figures S2-S3).

### ddPCR assays

Three separate ddPCR reactions were prepared for the detection of MTBC (duplex: *rpoB* and *IS6110*), HIV-1 (duplex: *pol* and *gag*) and human housekeeping gene (single-plex: *RPP30*) using a total of 1 µg total DNA each for MTBC and HIV-1 assays and 100 ng for the *RPP30* assay. The inclusion of two *Mtb* targets, *rpoB* and *IS6110*, enhances sensitivity and aids discrimination between high and low mycobacterial loads, characterized by double and single positive results, respectively.^19^ The ddPCR method applied for HIV-1 quantification was previously developed for use in blood-derived cells^25^, *RPP30* was included to quantify and normalise for human cell count variations across samples. DdPCR assay set-up, accuracy and calculation of MTBC ddPCR limit of detection (LOD) and representative ddPCR amplification plots for samples and no template controls are detailed in Supplementary Materials.

All pathogen ddPCR data are presented as either total copies/biopsy, or biopsy segment, and gene copy number per million human cells (copies/million cells), unless otherwise specified. Human cell number was calculated estimating two copies of *RPP30* DNA represents one human nucleated cell (see Supplementary Methods). Normalising to cell number allowed pathogen abundance comparison across varying sample types/sizes. *RPP30* copies/µg were compared between biopsy sample types for assessment of tissue cellularity and the relationship to pathogen abundance. For each sample, MTBC and HIV-1 ddPCR results were considered positive if DNA of either gene was detectable above background signal (i.e.: *rpoB* or *IS6110* and *gag* or *pol*, respectively).

### Statistics

Descriptive statistics included median values and interquartile range (IQR) for continuous variables and absolute number (n) and frequency (%) for categorical variables. To assess significant differences between groups, Fisher’s exact or Kruskal–Wallis tests and Wilcoxon rank sum tests were used for categorical and continuous variables, respectively. A p-value of <0.05 was considered statistically significant. Statistical analyses were conducted using GraphPad Prism Version 9.3.1. For the duplex *rpoB/*IS*6110* and *gag/pol* assays, the categorical variable was the proportion of patients in whom DNA of either gene was detected, and the continuous variable was the number of copies of *rpoB* and IS*6110*/million cells. Sample size was calculated for the original study ^17^ and samples used in this study represent a convivence sample, such that a formal power calculation was not performed.

## RESULTS

### Participant characteristics and total DNA samples for ddPCR derived from spinal biopsies

Of the 31 patients enrolled into the parent study, 25 had sufficient additional tissues collected for the ddPCR sub-study, 10 diagnosed *culture-confirmed STB*, 6 *Xpert Ultra-confirmed STB*, and 9 were diagnosed *Not STB* (Figure 1). Patient demographic, clinical and diagnostic characteristics are presented in Table 1 and Supplementary Tables S1-2. Half (8/16) of the patients with confirmed STB, and a third (3/9) of those diagnosed *Not* STB, were males. Those with confirmed SPTB were significantly younger than those without (median age: 40 years *culture-confirmed STB*, 42 years *Xpert-confirmed STB*, 53 years *Not STB*, p=0.02). Of the 25 patients, thirteen (52%) were also PLWH (100% ART established), with no difference in HIV status by TB diagnostic group (p=0.22). All reported ART adherence, yet seven (54%) had detectable peripheral VLs (median copies/ml: 446 (IQR: 294-676)), and 4/13 (31%) were virally suppressed (VL unavailable for 2). Seventeen (68%) patients had an open surgery large tissue biopsy specimen collected, whilst eight (32%) underwent percutaneous CT-guided needle biopsy, from which a single core biopsy specimen was collected.

**Figure 1.**
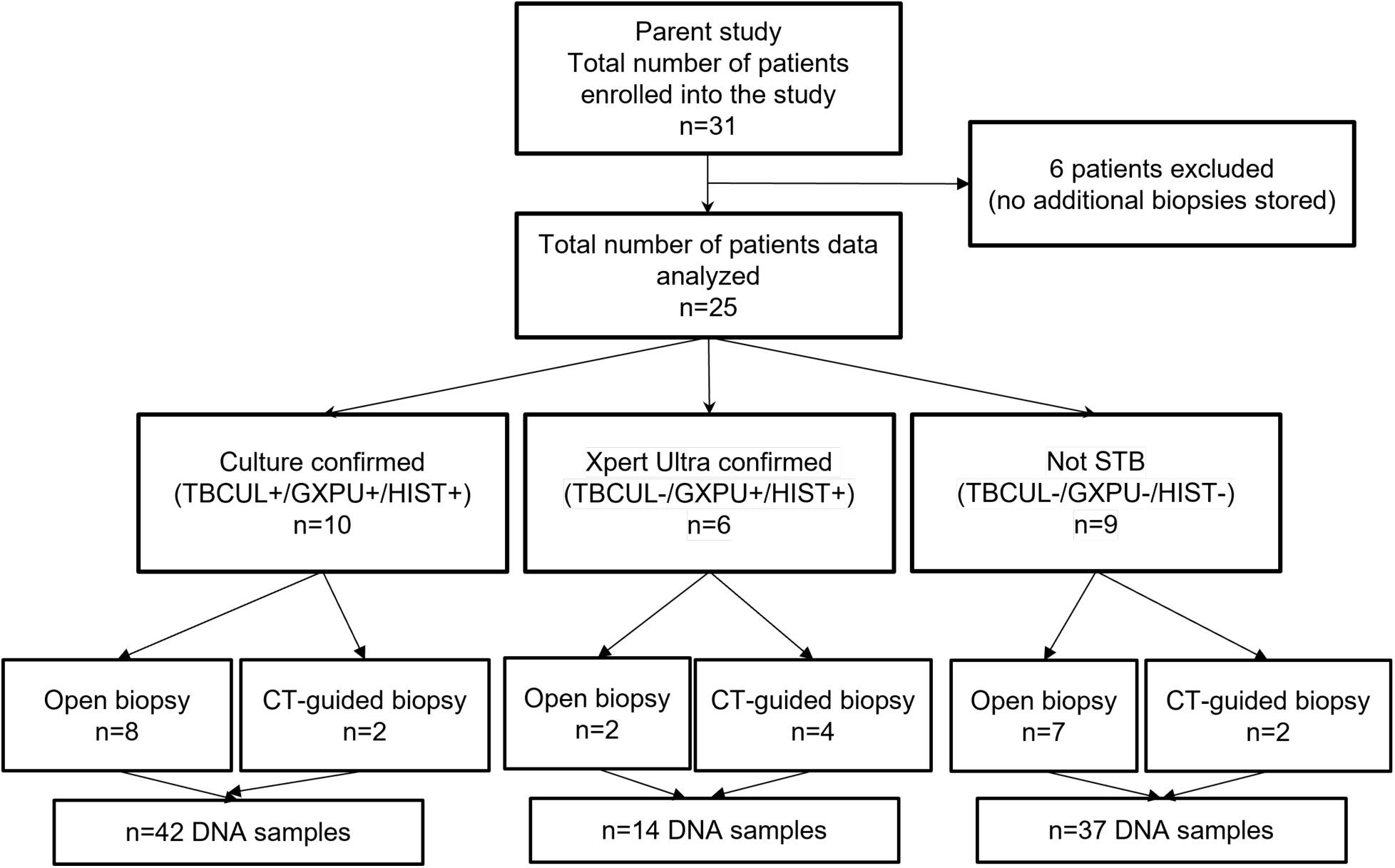
**Flow diagram of study patients including diagnostic, surgical, and ddPCR sample categories**. GXPU: Xpert MTB/RIF Ultra; TBCUL: MGIT TB culture; HIST: histology including histopathology and Ziehl-Neelsen stain for acid fast bacilli.

**Table 1.** Study patient characteristics, stratified by patient diagnostic group.

|  | All<br>Patients<br>(n=25) | TBCUL+<br>GXPU+HIST+<br>Culture<br>confirmed STB<br>(n=10) | TBCUL-<br>GXPU+HIST+<br>Xpert Ultra<br>confirmed STB<br>(n=6) | TBCUL-<br>GXPU-HIST-<br>Not STB<br>(n=9) |
| --- | --- | --- | --- | --- |
| Total biopsy segments | 93 (100%) | 42 (45%) | 14 (15%) | 37 (40%) |
| Age, years | 48 (36-53) | 40 (29-49) | 42 (38-44) | 53 (49-58) |
| Male sex | 11 (44%) | 5 (50%) | 3 (50%) | 3 (33%) |
| HIV-1 positive patients | 13 (52%) | 6 (60%) | 4 (67%) | 3 (33%) |
| Undetectable HIV-1 VL | 4/11<sup>A</sup> (36%) | 2/5<sup>A</sup> (40%) | 1/3<sup>A</sup> (33%) | 1 (33%) |
| Detected HIV-1 plasma VL | 446 (294-791) | 372 (292-391) | 1137 and 49959 | 60 and 676 |
| Previous TB history | 14 (56%) | 5 (50%) | 4 (67%) | 5 (56%) |
| TB therapy initiated before biopsy | 9 (28%) | 3 (30%) | 5 (83%) | 1 (11%) |
| TB culture positive | 10 (40%) | 10 (100%) | 0 (0%) | 0 (0%) |
| Xpert MTB/RIF Ultra + | 16 (64%) | 10 (100%) | 6 (100%) | 0 (0%) |
| Histopathology + | 12/19<sup>B</sup> (63%) | 7/7 (100%) | 5/5 (100%) | 0/7 (0%) |
| AFB + | 2/18<sup>C</sup> (11%) | 2/8 (25%) | 0/5 (0%) | 0/5 (0%) |
| Open surgery biopsy | 17 (68%) | 8 (80%) | 2 (33%) | 7 (78%) |
| CT-guided needle biopsy | 8 (32%) | 2 (20%) | 4 (67%) | 2 (22%) |

Data are proportion (n) and (%) or median (IQR). <sup>A</sup>n=2 unavailable viral load (VL) results (1 culture confirmed and 1 Xpert Ultra confirmed); n=4 lower than detectable limit (LDL); <sup>B</sup>No histology report available in n=6 patient (3 culture confirmed, 1 Xpert Ultra confirmed, 2 Not STB); <sup>C</sup>No Ziehl-Neelsen staining report for acid fast bacilli (AFB) available in n=7 patients (2 culture confirmed, 1 Xpert Ultra confirmed, 4 Not STB);. +: positive; -: negative; CT: Computed tomography; CUL: culture; GXPU: Xpert MTB/RIF Ultra; HIST: histology; TB: Tuberculosis.

From the 25 biopsies, DNA from 93 individual segments was analysed, with open biopsies portioned into five segments (n=17, 85 DNA samples) and CT-guided biopsies analysed as one segment/patient (n=8) (Table 1, Supplementary Figure S1). Open biopsies yielded approximately 100-fold more DNA and human cells (quantified as two *RRP30* copies/cell) than CT-guided biopsies (p<0.0001). There was no difference in the amount of DNA recovered or human cellularity based on open biopsy tissue type, STB diagnostic group or HIV-1 status (Supplementary Figure S4).

Of the 93 biopsy segments, 87 (94%) produced valid MTBC ddPCR data (>10,000 droplets), with 62 (67%) samples providing 1 µg of DNA for reactions (Supplementary Figures S5-S6, Table S3). The remaining 31 (33%) samples had less DNA, with a minimum of 46 ng added per reaction and 90 ng the lowest DNA amount that yielded a positive MTBC signal. MTBC assay validation using purified *Mtb* DNA determined detection limits of 2.29 *rpoB* copies/20µl (≈2 bacilli equivalent) and 9.67 IS6110 copies/20µl reaction (Supplementary Figures S5-S6).

### ddPCR has high sensitivity to detect MTBC DNA in biopsies from patients with current STB and previous PTB

First, to compare MTBC ddPCR performance against Xpert Ultra and TB culture, we combined results from the multiple segments obtained from each biopsy to analyse equivalent total biopsy results. MTBC DNA (*rpoB* or *IS6110*) was detected >LOD in 19/25 (76%) total biopsy samples of suspected STB patients: 10/10 (100%) *culture-confirmed* patients, 5/6 (83%) *Xpert Ultra-confirmed* patients and 4/9 (44%) *Not STB* patients (Table 2, individual biopsy results in Supplementary Table S2). The one negative Xpert Ultra-confirmed sample was a CT-guided needle biopsy from a patient initiated on TB treatment before biopsy. The four MTBC DNA patients diagnosed *Not STB* all had previous PTB and one also lymph node (LN) TB; two were *rpoB+IS6110+* and two *rpoB+IS6110-*. Two of these four patients subsequently died (Figure 2A, Supplementary Tables S1-S2).

**Figure 2.**
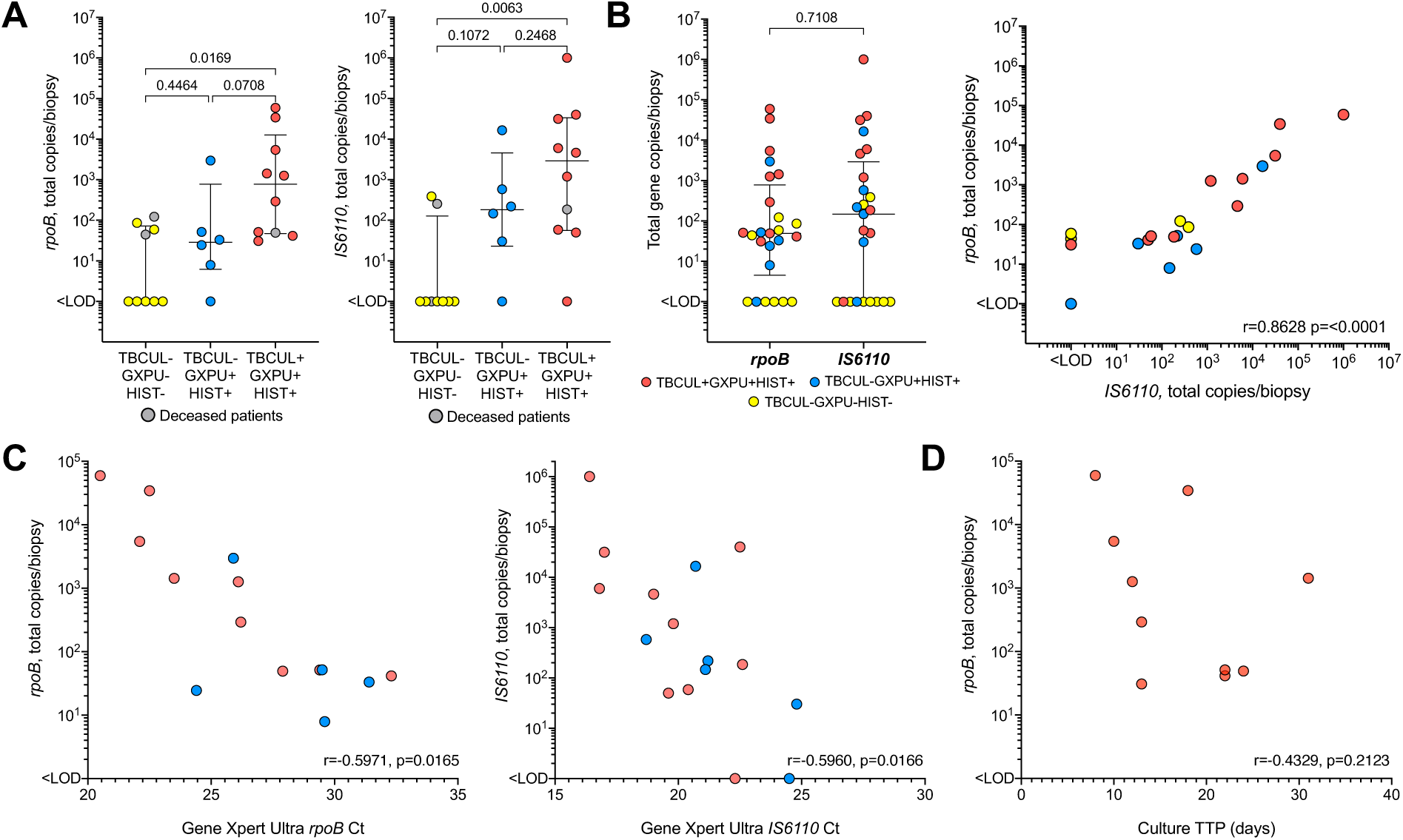
Total biopsy ddPCR MTBC gene copies from adult patients investigated for STB, undergoing either percutaneous CT-guided needle biopsy or open biopsy in theatre. **A**) Total *rpoB* and *IS6110* copies per biopsy, stratified by STB diagnostic group. Grey dots represent deceased patients who passed away after diagnosis. **B**) Total *rpoB* and *IS6110* copies, per total biopsy, for all patients combined. Data plotted as median + IQR and the Spearman’s correlation between *IS6110* and *rpoB* total copies per biopsy. **C**) Total *rpoB* ddPCR copies and *IS6110* copies per biopsy, correlated with Gene Xpert MTB/RIF Ultra (GXPU) *rpoB* and *IS6110* Ct values, respectively, colored by STB diagnostic group in A-B. One GXPU positive did not have *rpoB* Ct values. **D**) Total *rpoB* copies/biopsy correlated with MGIT TB culture (TBCUL) time to positivity (TTP). Dots represent combined biopsy segment results for each patient. (A-B) Kruskal-Wallis test with Benjamini, Krieger and Yekutieli FDR (q-value) and (B-D) Spearman correlation coefficients and p values calculated using Spearman’s test. HIST: Histology, LOD: Limit of detection.

MTBC total gene copies/biopsy ranged 8-59,144 for single-copy *rpoB* and 30-1,001,775 for multi-copy *IS6110*, with *rpoB* and *IS6110* total copies/biopsy positively correlated (r=0.8628, p<0.0001, Figure 2B). There was a significant difference in total gene copies/biopsy for both *rpoB* and *IS6110* between those diagnosed as *Not STB* with *culture-confirmed*, but not *Xpert Ultra-confirmed* STB patients (Figure 2A). Total copies/biopsy of *rpoB* and *IS6110* were significantly inversely correlated with Xpert Ultra *rpoB* and IS6110 Ct values (r≤-0.597, p<0.017), but ddPCR *rpoB* copies (i.e. bacilli equivalents) were not correlated with culture time to positivity (Figure 2C-D).

### MTBC DNA detection and abundance varies between segments of the same biopsy

To investigate the spatial variation in *Mtb* presence and abundance within biopsy tissues, we next compared results between individual biopsy segments (see Figure 3A and Supplementary Figures S2-S3). 45/87 (52%) biopsy segments with valid MTBC results had detectable levels of *IS6110/rpoB*: 29/87 (34%) *rpoB+IS6110+*, 16/87 (18%) single-positive for *rpoB* (15) or *IS6110* (1) (Table 2). Notably, 10/12 *rpoB* single-positive segments came from biopsies that also had double-positive segments, with the single-positive segments generally having ≤20 μg DNA extracted (Figure 3B). The greater detection of *rpoB* single-positives, compared to IS6110, may be explained by the brighter fluorophore used for this probe increasing sensitivity to detect positive droplets above the fluorescence threshold for samples with lower DNA content (Supplementary Figures S7-9). IS6110-negative *Mtb* clinical strains also exist.^28^

**Figure 3.**
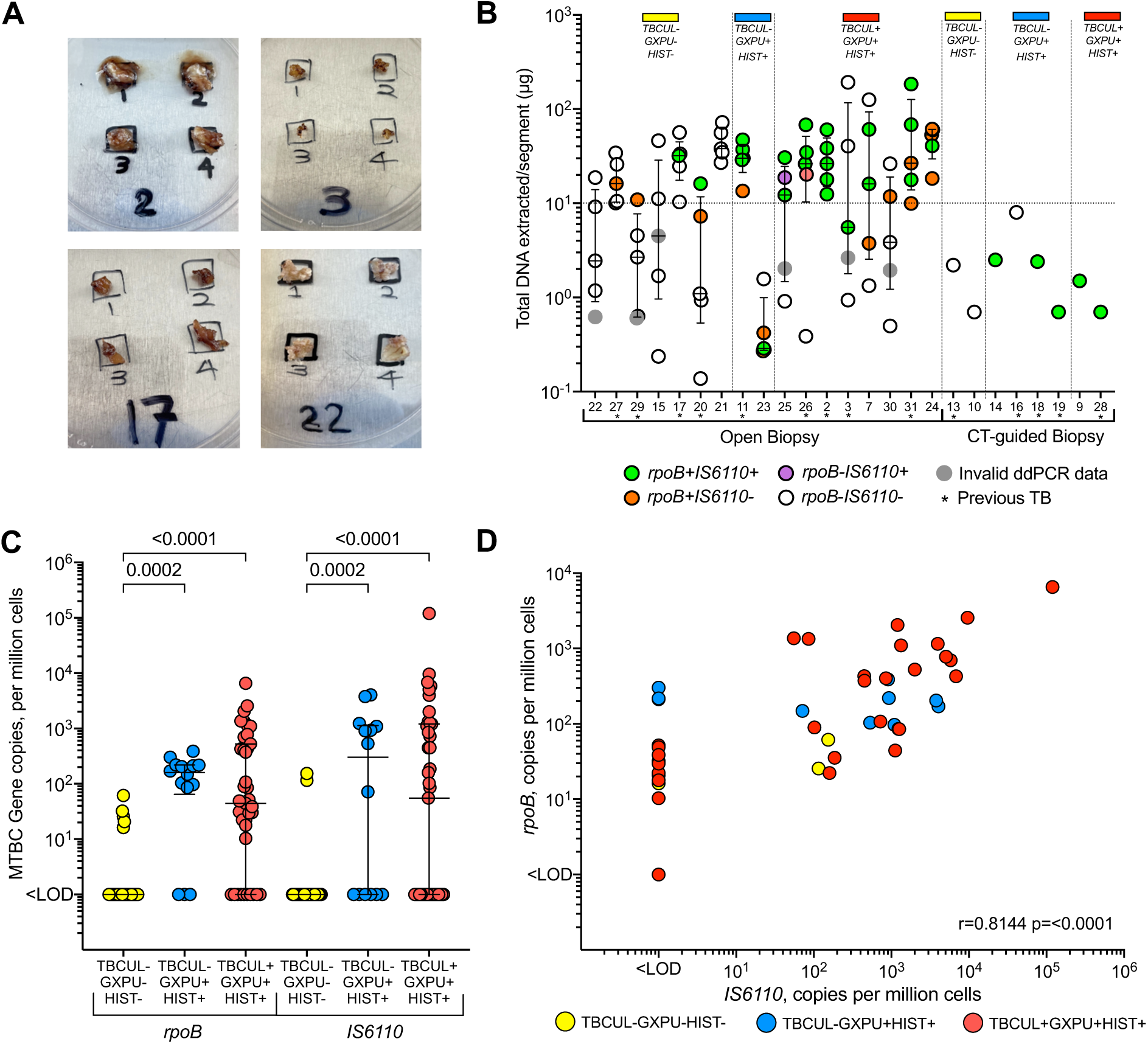
**MTBC *rpoB* and *IS6110* copies detected by ddPCR**. **A)** Representative images of segmented open spinal biopsies with diseased soft (##2,3,17) or hard bone (#22), B) MTBC DNA detection in total DNA extracted per segment of open biopsy patients, or some the entire CT-guided needle biopsy core, stratified by STB diagnostic group. Individual dots represent total DNA extracted per biopsy segment (n=5 open biopsy, n=1 CT-guided biopsy). Double positives (*rpoB+IS6110*+, green dots; no *rpoB* or *IS6110* detected (*rpoB-IS6110-,* white dots with black border); single positives (*rpoB+IS6110-,* orange dots; single positives (*rpoB-IS6110+,* purple dots). X-axis labelled with patient ID. **C**) Gene copies in biopsy segments, stratified by STB diagnostic grouping. *rpoB* and *IS6110* copy numbers per million human cells for n=87 segments with valid ddPCR data. <LOD was ‘undetectable’. Kruskal-Wallis Test with Benjamini, Krieger and Yekutieli FDR (q-value). Lines show median values with IQR. **D)** *IS6110* versus *rpoB* copies/million cells for the 87 biopsy segments with valid ddPCR data. X and Y axes are log10. Spearman correlation coefficients and p values were calculated using Spearman’s test. GXPU: Xpert MTB/RIF Ultra, HIST: Histology, LOD: Limit of detection, TBCUL: MGIT TB culture.

MTBC DNA was detected in 29/39 (74%) segments from *culture-confirmed* and 11/14 (79%) segments from *Xpert Ultra-confirmed* STB patients. 5/34 (15%) *Not STB* segments had detectable *rpoB*/*IS6110*, 2/34 (6%) *rpoB+IS6110+* (two separate patients) and one *Not STB* patient with previous PTB who died had 2 positive segments (Figure 3B, Table 2).

MTBC gene copies/million cells were significantly lower in segments from patients diagnosed *Not STB* compared to STB (p≤0.0002) (Figure 3C). For *culture-confirmed* patients, copies/million cells ranged 10-6,558 for *rpoB* and 55-119,698 for *IS6110*. For *Xpert Ultra-confirmed* patients, these were 86-389 and 72-4,045, respectively. In comparison, the five positive biopsy segments from the *Not STB* patients, had 16-61 and 115-154 copies/million cells, respectively (Figure 3C). Combining samples from all patients, *IS6110* and *rpoB* copies/million cells were positively correlated (r=0.814, p<0.0001, Figure 3D). However, when analysing by STB category, correlations remained for *culture-confirmed* (r=0.8265, p<0.0001) and *Not STB* patients (r=0.642, p<0.0001), but not *Xpert Ultra-confirmed* patients (r=0.1264, p=0.6637). Notably, most *Xpert Ultra-confirmed* patients were initiated on TB therapy before biopsy (Supplementary Table 2), thus more extracellular DNA could have been detected.

### HIV-1 DNA is detectable in spinal biopsy tissue from PLWH, irrespective of STB diagnosis and peripheral viral load

Next, we investigated the ability of ddPCR to detect HIV-1 DNA in spinal biopsy tissues from the 13 PLWH, detecting HIV-1 DNA (*pol*/*gag*) in all 13 biopsies. 34/37 (92%) tissue segments had detectable HIV-1 DNA, 91-100% across STB diagnostic group: 22/37 (60%) were *pol+gag+,* 10/37 (27%) were *pol-gag+*, and 2/37 (5%) were *pol+gag-* (Table 2). Similar to the results for MTBC DNA, double-positive open biopsy segments mostly had >10µg DNA extracted, reflecting high tissue cellularity (Figure 3A and 4A). Of the three patients with HIV-1 DNA detected in less than 5 open biopsy segments, two had lower than detectable (LDL) blood VL, and one had VL 60 copies/ml. In patients who underwent CT-guided biopsies two had LDL VL, 4/6 (67%) were *pol+gag+,* 1/6 (17%) *pol-gag+*, and 1/6 (17%) *pol-gag+* (Figure 4A).

**Figure 4.**
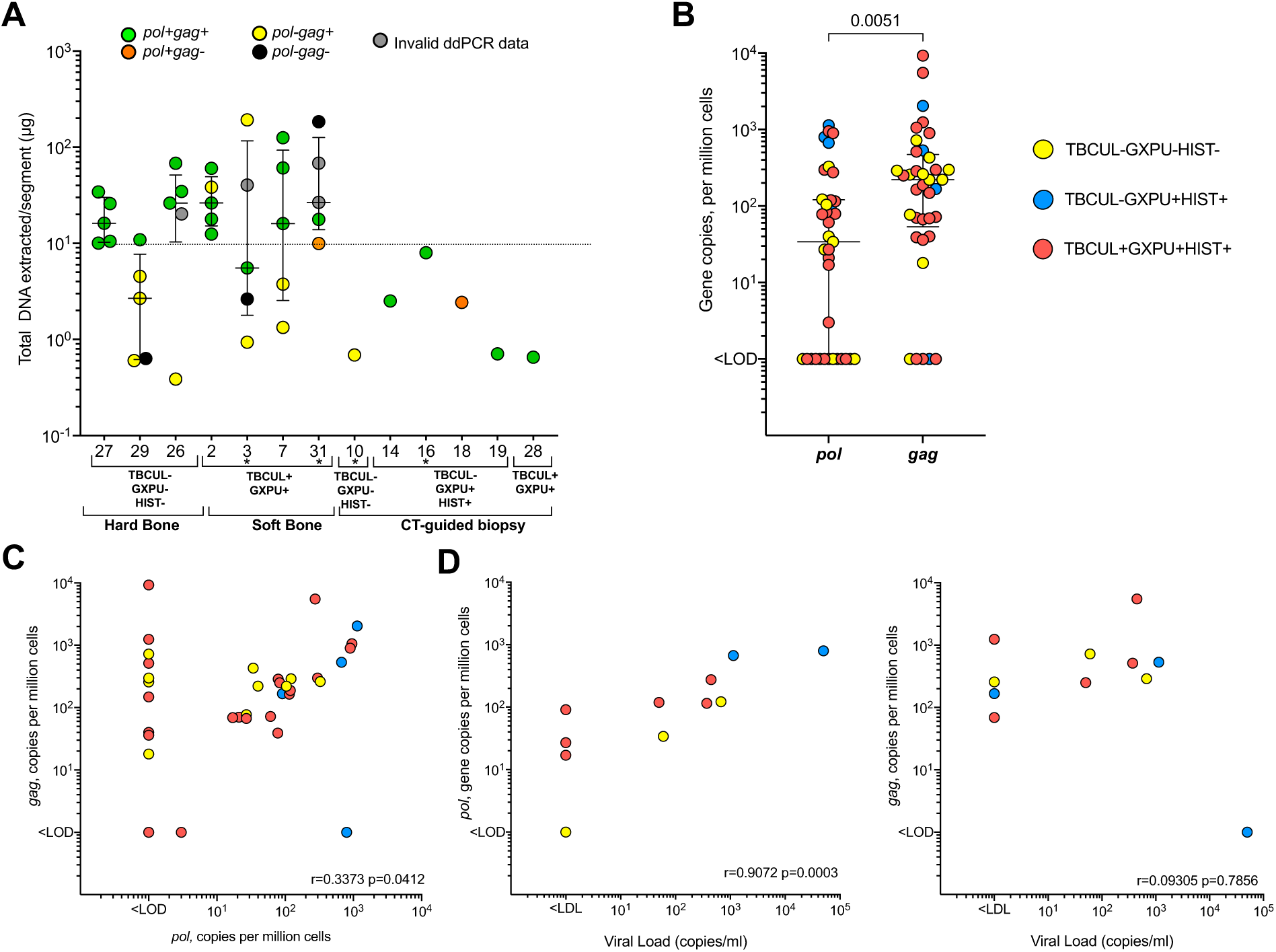
Detection of HIV-1 DNA in spinal biopsy tissue from PLWH and the relationship to blood viral load. **A**) HIV-1 DNA detection in total DNA extracted per segment of open biopsy from PLWH, stratified by STB diagnostic group and type of tissue extracted during surgery. Individual dots represent total DNA extracted per five biopsy segments from each patient. X axis is patient ID. *Pol* and *gag*: HIV-1 genes. Invalid ddPCR: samples with droplet counts <10 000, no detection by software. * on patient ID indicates those with blood viral load lower than detectable levels. **B**) Detection and **C**) correlation of HIV-1 *pol* and *gag* gene copies per million cells in DNA from biopsy tissues of adult suspected spinal tuberculosis patients. **D**) Correlation between spinal biopsy maximum detected *pol* or *gag*, copies per million cells across tested biopsy segments and plasma HIV-1 viral load. Correlation coefficients and p-values were calculated with Spearman’s test. n=2 patients had unknown viral loads and were excluded from D and E analysis*. GXPU: Xpert MTB/RIF Ultra; HIST: Histology; TBCUL: MGIT TB culture*.

Across all PLWH segments, *gag* copies/million cells were significantly higher than *pol*, median (IQR) 254 (75-519) *vs.* 97 (39-305), respectively (p=0.0051), and only modestly correlated (r=0.33, p=0.04) (Figure 4B-C). However, restricting the analysis to *pol+gag+* segments (potentially representing a higher proportion of intact genomes, although not directly measurable by the assay), the positive correlation improved (r=0.7698, p<0.0001).

There was also a strong positive correlation between HIV-1 blood VL and the maximum segment *pol* copies/million cells for each patient (r=0.9072, p=0.0003), but not for *gag* (r=0.09305, p=0.7856, Figure 4D). This may support *gag* single positive segments being indicative of defective virus, with *gag* being closer to the 5’ end of the HIV-1 genome.

### MTBC and HIV-1 abundance are increased in spinal tissue segments where both pathogens are detected

To investigate the variation in distribution of both pathogens across biopsy tissues, we first compared the number of open biopsy segments in which each pathogen was detected, finding HIV-1 more widely distributed than *Mtb*: HIV-1 DNA (*gag*/*pol*) detected in 28/31 (90%) segments from PLWH, and MTBC DNA detected in only 35/47 (74%) segments from confirmed STB patients (Figure 3A and 4A). Analysing the impact of HIV-1 co-infection on MTBC (*rpoB/IS6110*) detection in total biopsies, we found no significant difference by patient HIV-1 status (Supplementary Table S4). There was also no difference by HIV-1 status in *rpoB* or *IS6110* total DNA copies/biopsy (Figure 5A), in line with our previous findings based on *Mtb* abundance estimated from culture and Xpert Ultra^17^. However, when analysing biopsy segment results normalised to tissue human cellularity, *rpoB* and *IS6110* copies/million cells were significantly higher in biopsies from PLWH (p≤0.01, Figure 5A).

**Figure 5.**
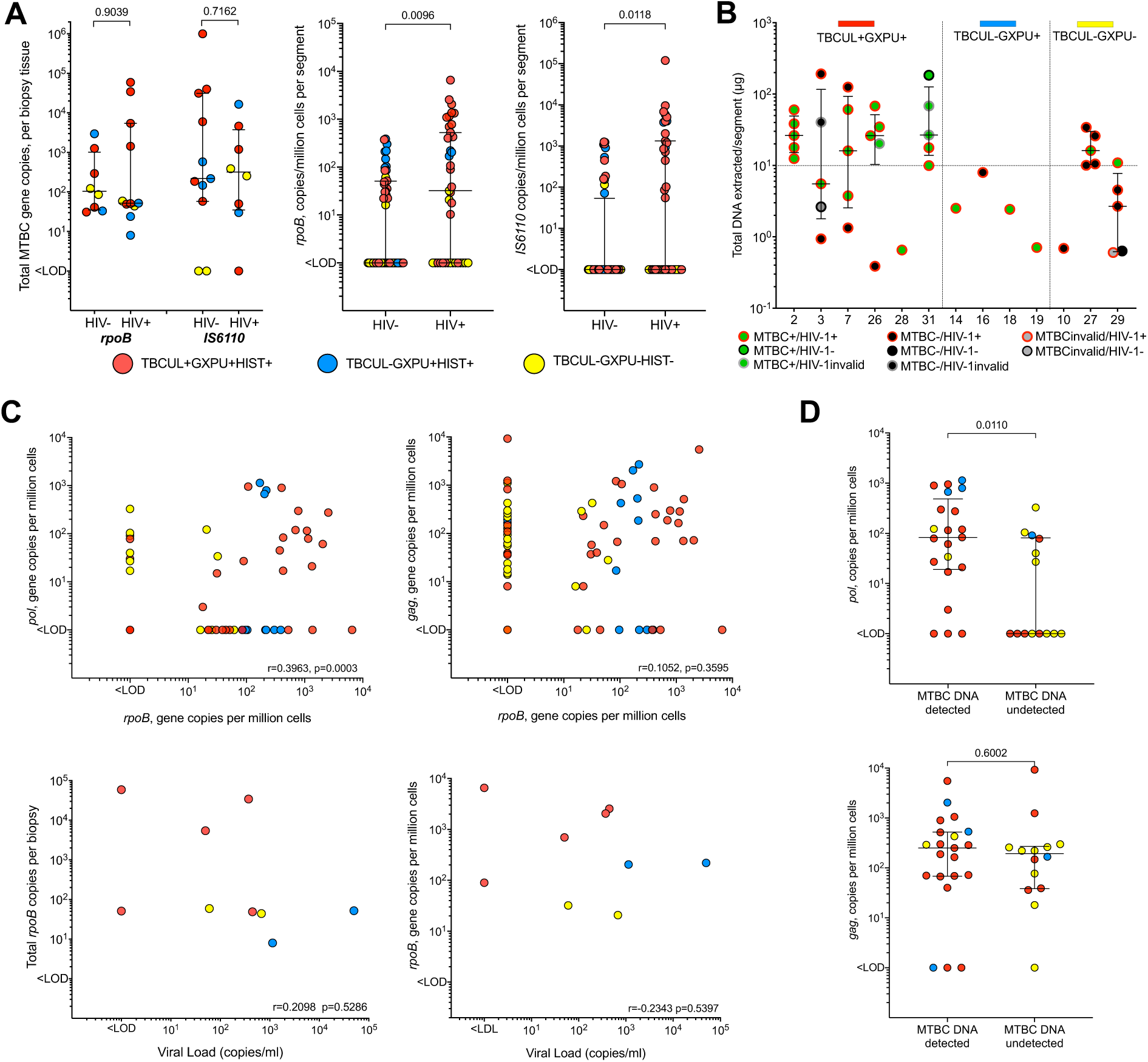
Relationship between MTBC DNA abundance and HIV-1-status, gene copies, and co-detection of MTBC DNA in spinal TB biopsy tissues. **A)** Total MTBC *rpoB* and *IS6110* copies/biopsy tissue, by HIV status, and *rpoB* and *IS6110* copies/million cells per biopsy segment, by HIV status. (**B)** Detection of both MTBC and HIV-1 DNA in biopsy segments from open and CT-guided biopsy from PLWH, stratified by STB diagnostic group. Individual dots represent total DNA extracted per biopsy segment for either open (n=5/patient) or CT-guided biopsies (n=1/patient). MTBC+ = detection of *rpoB* or *IS6110* and HIV-1*+* = detection of *pol or gag.* X axis is MUTI patient ID. MUTI016 was on empiric anti-tuberculous therapy at the time for concurrent pulmonary TB with undetectable MTBC DNA yet detectable HIV-1 DNA. (**C**) Correlation between HIV-1 *pol* or *gag* copies/million cells with *rpoB* copies/million cells in each biopsy segment and correlation between patient HIV-1 VL in the blood and spinal biopsy total detected *rpoB* copies or maximum biopsy segment *rpoB* copies/million cells. (**D**) HIV-1 gene copies/million cells for *pol* and *gag* in biopsy samples stratified by whether MTBC DNA (either *rpoB or IS6110*) was detected by ddPCR. P values were calculated with Mann-Whitney tests (A,D). Correlation coefficients and p-values were calculated with Spearman’s test (C)). Lines show median values with interquartile range (IQR). Dots in all graphs coloured by STB diagnostic group, as per (A). *GXPU: Xpert MTB/RIF Ultra; MTBC: Mycobacterium tuberculosis complex; TBCUL: MGIT TB culture. LOD: limit of detection*.

To investigate whether there exists a synergistic interaction between HIV-1 and *Mtb* in spinal tissues we next compared their presence and abundance within 35 biopsy segments from the 13 PLWH (STB and Not STB) with valid ddPCR data for both MTBC and HIV-1 DNA. 11/13 (92%) patients had MTBC and HIV-1 DNA co-detected in at least one of their open biopsy segments (n=7) or CT-guided biopsy (n=4) (Figure 5B). 20/35 (57%) segments had both MTBC and HIV-1 DNA detected (*rpoB/IS6110* AND *pol*/*gag*), of which 70% (14/20) were *rpoB+IS6110+* and *pol+gag+,* 10% (2/20) *rpoB+IS6110+* and *pol+/gag+,* and 10*% rpoB+IS6110-* and *pol+gag+*.

Within individual biopsy segments, MTBC single copy gene *rpoB* copies/million cells were positively correlated with HIV-1 *pol* DNA copies/million cells (r=0.3963, p=0.0003) in the same biopsy segment, but not HIV-1 *gag* copies/million cells (p=0.3595) (Figure 5C). However, maximum biopsy segment *rpoB* copies/million cells and total biopsy MTBC *rpoB* copies were not significantly correlated with blood VL. Therefore indicating *Mtb* abundance at STB tissue sites is positively correlated with local HIV-1 viral genome abundance and not peripheral VL (Figure 5C). Finally, testing the impact of co-localised MTBC presence on tissue segment HIV-1 abundance, we found *pol* copies/million cells were significantly higher in biopsy segments with MTBC DNA co-detected (p=0.0110), whilst there was no significant difference for *gag* copies/million cells (p=0.6002) (Figure 5D).

## DISCUSSION

Here we report the findings of the first study, to our knowledge, to test for MTBC DNA and HIV-1 DNA in spinal biopsy tissue from patients investigated for STB, living in a setting with a high incidence of both TB and HIV-1 infection^29^. The ddPCR assays successfully detected and quantified MTBC DNA in biopsy tissues from 94% STB confirmed patients and HIV-1 DNA in 100% of biopsies from ART-established PLWH, including four peripherally virally suppressed. The ddPCR assay also detected MTBC DNA in biopsies from four patients in which *Mtb* could not be detected by culture or Xpert Ultra, all of whom had a previous PTB diagnosis. Cutting biopsies into segments and normalising pathogen abundance to the localised human cell number enabled investigation of the tissue distribution and co-localised interaction between *Mtb* and HIV-1. Notably, HIV-1 was more widely distributed, being detected in 91% of segments, compared to 74% for *Mtb*. Importantly, whilst maximum tissue localised HIV-1 *pol* copies/million cells strongly correlated with blood VL, MTBC *rpoB* copies/million cells also positively correlated with matched tissue segment HIV-1 *pol* copies/million cells and not blood VL. Thus, our findings support a reciprocal tissue segment localised relationship between *Mtb* and HIV-1 abundance. This finding opens the door for future studies to characterise the immunological difference between co-and mono-infected tissue segments as a means to more precisely characterise the co-infection immune microenvironment and its role in enhancing HIV-1/*Mtb* reservoir persistence/expansion.

In addition to our creative approach of cutting biopsies into multiple segments to investigate the local microenvironment interaction between *Mtb* and HIV-1 we also normalised pathogen abundance to tissue segment human cell count by measuring copies of *RRP30*, unlike other recent studies which have used ddPCR to investigate MTBC-DNA in blood^22,30^. This enabled identification of a significantly higher abundance of *Mtb* at the site of STB in PLWH, that was not previously found when merely comparing *Mtb* abundance based on total biopsy culture or Xpert values.^17^ We had also postulated that with *Mtb* infection, the cellular composition of tissues at the site of disease may be more abundant with recruited and infiltrated cells, however, neither the abundance of *Mtb* infection in the spine nor the abundance of HIV-1 co-infection was associated with biopsy cellularity, although there were trends observed.

The high sensitivity of ddPCR, enabled low-level detection of MTBC DNA in four patients who were clinically diagnosed STB negative based on culture and Xpert Ultra, all of whom had previous PTB and one also LN TB, two being PLWH. Notably, 68% of the patients with *Mtb* DNA detected in their biopsies by ddPCR had a previous TB episode. *Mtb* may have seeded in the spine via a haematogenous route following pulmonary or LN disease. In the case of the four patients diagnosed *Not STB,* the detected MTBC DNA may have arisen from this previous haematogenous seeding of *Mtb* in the spine that was no longer viable so culture negative, or it represents a misdiagnosis by culture and Xpert Ultra. Positivity was unlikely due to cross-contamination or non-specific amplification, as all 14 biopsy segments from those diagnosed *Not STB* with no previous TB were *rpoB*-*IS6110-,* as were all negative controls.

Identifying tissue-based and cellular reservoirs of HIV-1 in ART-treated patients is crucial for finding a cure. Studies have found HIV-1 DNA in bone marrow cells^31,32^ and higher than expected levels of HIV proviral DNA in postmortem spinal cord tissue^33^. Our detection of HIV-1 DNA in biopsies from patients with no detectable viral loads in the blood, potentially indicate tissue-specific HIV-1 presence and persistence that is enhanced by *Mtb* tissue presence. As we’ve previously reviewed^12^, the inflammatory state during active *Mtb* infection has the potential to both enhance viral replication, by activating HIV-1 gene transcription from the HIV long terminal repeat promoter region, and creating an expanded cellular niche that is susceptible to HIV-1 infection. HIV-1 establishes latency shortly after infection through integrating into the host genome as a provirus.^34^ As ART only inhibits actively replicating virus, it cannot eliminate the virus from latently infected cells, which persist through mechanisms such as homeostatic proliferation^35^ and represent a source of viral rebound, if treatment is interrupted.^36^

Since the HIV reservoir is composed of predominantly replication defective virus, the poor correlation and higher detection of *gag* verses *pol* copies we detected could reflect defective proviral genomes that are hypermutated or have large deletions^37^. Alternatively, this could be due to varying primer sensitivity or off-target amplification. Future studies could include the Intact Proviral DNA Assay (IPDA), a similar ddPCR method with different targets that have increased statistical likelihood of representing intact genomes^24^ and combining ddPCR with flow cytometry or single-cell omics to better characterize the HIV-1 reservoir cells.

The small sample size and single hospital site are limitations of our study, primarily due to the COVID-19 pandemic’s impact changing patients’ hospital seeking nature. Future work should assess *Mtb* viability through detecting MTBC *16S* mRNA alongside *rpoB* and *IS6110* DNA detection. Our study also highlights the potential improved sensitivity and quantitative accuracy of ddPCR for STB diagnosis, however assessment on larger sample sizes and more diverse patient groups is required to clarify ddPCR’s diagnostic performance for STB. While ddPCR’s current use is limited by cost and infrastructure^38,39^, recent chip-based innovations may overcome these barriers. Further studies should also compare MTBC and HIV-1 DNA detection in peripheral blood compared to disease sites and how these change with treatment, though this would require repeated biopsies. Finally, we did not measure HIV RNA or proteins in tissue biopsies, limiting our ability to determine local HIV replication at MTBC colocalization sites.

In conclusion, ddPCR provides highly sensitive detection of *Mtb* in STB biopsies, including in the context of HIV-1 infection, with potential superior sensitivity over Xpert Ultra, and the added benefit of absolute quantification. Through segmenting biopsies into multiple pieces we confirmed a tissue co-localised relationship between *Mtb* and HIV-1 abundance, suggesting both pathogens facilitate expansion of the other in spinal sites of co-infection. This needs to be considered with detection methods and tailored treatment strategies that ensure penetrance into anatomical sites such as the spine. The contribution of the TB microenvironment as both a reservoir and site of HIV reservoir activation requires further investigation.

## DATA SHARING

Deidentified data from this study will be made available upon request with investigator support, after approval of a proposal, and with a signed data access agreement. For data access, please contact the corresponding author.

## ACKNOWLEDGMENTS.

We thank Nomsa Yekiso, Nosipho Mncwabe, Dr Qonita Hartley, Dr Chad Centner, Ms Nawaal Adikary and Zeenat Hoosen, orthopaedic surgical teams and our patients for helping, participating and contributing to global evidence. The following reagent was obtained through the NIH HIV Reagent Program, Division of AIDS, NIAID, NIH: HIV-1 Lymphadenopathy-Associated Virus (LAV)-Infected 8E5 Cells, ARP-95, contributed by Dr Thomas Folks. The works was supported by Wellcome (203135Z/16/Z); Academy of Science of SA (ASSAf); ORU, UCT; UCT Merit Award. For the purposes of open access, the authors have applied a CC-BY public copyright to any author-accepted manuscript arising from this submission.

## CONTRIBUTORS

RW collected tissues, performed assays, analysed data and drafted the manuscript. ACK, MRA, MH, and ARM contributed to study design, supported data collection and analysis, and edited the manuscript. RND and ML aided study design, input and sample collection.

---

**Table S2.** Total biopsy and biopsy segment ddPCR detection of MTBC and HIV-1 DNA, stratified by STB diagnostic group.

|  | TBCUL+GXPU+<br>HIST+ Culture<br>Confirmed STB | TBCUL-GXPU+<br>HIST+ Xpert Ultra<br>Confirmed STB | TBCUL-GXPU-<br>HIST-<br>Not STB | All Patients |
| --- | --- | --- | --- | --- |
| **Total Biopsy, all patients** | **n=10** | **n=6** | **n=9\*** | **n=25** |
| MTBC DNA ddPCR+ | 10 (100%) | 5 (83%) | 4 (44%) | 19 (76%) |
| *rpoB*+*IS6110*+ | 9 (90%) | 5 (83%) | 2 (22%) | 16 (64%) |
| *rpoB*+*IS6110*- | 1 (10%) | 0 (0%) | 2 (22%) | 3 (12%) |
| *rpoB*-*IS6110*+ | 0 (0%) | 0 (0%) | 0 (0%) | 0 (0%) |
| *rpoB*-*IS6110*- | 0 (0%) | 1 (17%)<sup>#</sup> | 5 (56%) | 6 (24%) |
| **Biopsy segments, all patients** | **n=39** | **n=14** | **n=34** | **n=87** |
| MTBC DNA ddPCR+ | 29 (74%) | 11 (79%) | 5 (15%) | 45 (52%) |
| *rpoB*+*IS6110*+ | 19 (49%) | 8 (58%) | 2 (6%) | 29 (34%) |
| *rpoB*+*IS6110*- | 9 (23%) | 3 (21%) | 3 (9%) | 15 (17%) |
| *rpoB*-*IS6110*+ | 1 (3%) | 0 (0%) | 0 (0%) | 1 (1%) |
| *rpoB*-*IS6110*- | 10 (25%) | 3 (21%) | 29 (85%) | 42 (48%) |
| **Total biopsy, HIV+ patients** | **n=6** | **n=4<sup>^</sup>** | **n=3** | **n=13** |
| HIV-1 DNA ddPCR+ | 6 (100%) | 4 (100%) | 3 (100%) | 13 (100%) |
| *pol*+*gag*+ | 6 (100%) | 3 (75%) | 2 (67%) | 11 (84%) |
| *pol*+*gag*- | 0 (0%) | 1 (25%) | 0 (0%) | 1 (8%) |
| *pol*-*gag*+ | 0 (0%) | 0 (0%) | 1 (33%) | 1 (8%) |
| *pol*-*gag*- | 0 (0%) | 0 (0%) | 0 (0%) | 0 (0%) |
| **Biopsy segments, HIV+ patients** | **n=22** | **n=4<sup>^</sup>** | **n=11** | **n=37** |
| HIV-1 DNA ddPCR+ | 20 (91%) | 4 (100%) | 10 (91%) | 34 (92%) |
| *pol*+*gag*+ | 13 (59%) | 3 (75%) | 6 (55%) | 22 (59%) |
| *pol*+*gag*- | 1 (5%) | 1 (25%) | 0 (0%) | 2 (5%) |
| *pol*-*gag*+ | 6 (27%) | 0 (0%) | 4 (36%) | 10 (27%) |
| *pol*-*gag*- | 2 (9%) | 0 (0%) | 1 (9%) | 3 (8%) |

Data are presented as n (%) per diagnostic group. GXPU: Xpert MTB/RIF Ultra, HIST: Histology, TBCUL: MGIT TB culture. 87/93 biopsy segments had valid MTBC ddPCR data points and 37/41 HIV+ biopsy segments had valid HIV-1 ddPCR >10,000 droplets.
\*5/9 diagnosed *Not STB* had previous pulmonary TB, 4/5 of these were MTBC DNA ddPCR+.
<sup>#</sup>Sample was CT-guided biopsy and patient initiated on TB treatment before biopsy.
<sup>^</sup>All four CT-guided biopsies.

