## Supplementary for "Digital PCR detection of *Mycobacterium tuberculosis* and HIV-1 co-localization in spinal tuberculosis biopsies"

### TABLE OF CONTENTS

**SUPPLEMENTAL METHODS** 4

SEX AS A BIOLOGICAL VARIABLE 4
INCLUSION AND EXCLUSION CRITERIA SIGNS AND SYMPTOMS OF ADULT PATIENTS PRESENTING TO OUR UNIT FOR DIAGNOSIS.[1] 4
OPTIMISED DNA EXTRACTION METHOD FROM OPEN AND CT-GUIDED BIOPSIES 4
DROPLET DIGITAL PCR FOR THE DETECTION OF TB, HIV-1 AND RPP30 IN SPINAL BIOPSY SAMPLES. 5
ACCURACY AND CALCULATION OF LIMIT OF DETECTION OF MTBC DNA DDPCR ASSAY 6

**SUPPLEMENTAL TABLES** 7

TABLE S1. HISTOPATHOLOGICAL OR OTHER DIAGNOSIS FOR THE NINE PATIENTS NOT DIAGNOSED WITH STB. 7
TABLE S2. PER PATIENT DETECTION OF *MTB*, HIV-1 AND HUMAN DNA COPIES IN SPINAL TISSUES, STRATIFIED BY STB DEFINITION AND HIV STATUS. 8
TABLE S3. MEAN DROPLET COUNT AND MEAN TOTAL VOLUME OF DROPLETS FOR EACH DDPCR ASSAY 9
TABLE S4. DETECTION OF MTBC DNA BY PATIENT HIV-1 STATUS. 9
TABLE S5. OLIGONUCLEOTIDE INFORMATION, MTBC DNA DDPCR ASSAYS. 9
TABLE S6. OLIGONUCLEOTIDE INFORMATION FOR HIV-1 AND RPP30 DDPCR ASSAYS. 10
TABLE S7. REACTION SETUP PER DDPCR ASSAY (MTBC, HIV-1 AND RPP30 DETECTION) 11
TABLE 8. THERMAL CYCLING CONDITIONS, DDPCR ASSAYS 11

**SUPPLEMENTAL FIGURES** 12

FIGURE S1. TOTAL DNA EXTRACTION FROM SPINAL BIOPSY SPECIMENS. 12
FIGURE S2. REPRESENTATIVE IMAGES OF SEGMENTS OF OPEN SPINAL BIOPSIES WITH DISEASED HARD BONE (WHITE), SOFT BONE (MAROON) AND OTHER DISEASED TISSUE (MUTI024 AND MUTI023) (N=17). 13
FIGURE S3. REPRESENTATIVE IMAGES OF PERCUTANEOUS CT-GUIDED CORE BIOPSIES (N=8). 14
FIGURE S4. TOTAL DNA EXTRACTED AND CELLULARITY OF EACH PATIENTS' SPINAL BIOPSY TISSUE, STRATIFIED BY BIOPSY METHOD, TISSUE TYPE, SPINAL TB DIAGNOSIS AND HIV STATUS. 15
FIGURE S5. MTBC GENE COPY NUMBER PER 20μL WELL FOR A DILUTION SERIES OF H37RV MTB DNA. 16
FIGURE S6. REPRESENTATIVE QUANTASOFT DDPCR AMPLIFICATION PLOTS FROM MTB H37RV DNA DILUTION SERIES. 17
FIGURE S7. PROBABLE AND DEFINITE STB SPINAL TUBERCULOSIS PATIENTS MTBC DDPCR REPRESENTATIVE QUANTASOFT AMPLIFICATION PLOTS WITH MULTIPLE POSITIVE DROPLETS FOR TOTAL DNA SAMPLES EXTRACTED FROM SPINAL BIOPSY TISSUES. 18
FIGURE S8. NOT STB (TBCUL-GXPU-HIST-) PATIENTS (WITH PREVIOUS TB HISTORY) MTBC DDPCR REPRESENTATIVE QUANTASOFT AMPLIFICATION PLOTS WITH MULTIPLE POSITIVE DROPLETS FOR TOTAL DNA SAMPLES EXTRACTED FROM SPINAL BIOPSY TISSUES. 19

FIGURE S9. NOT STB (TBCUL-GXPU-HIST-) PATIENTS (NO PREVIOUS TB HISTORY) MTBC DDPCR REPRESENTATIVE QUANTASOFT AMPLIFICATION PLOTS FOR TOTAL DNA SAMPLES EXTRACTED FROM SPINAL BIOPSY TISSUES WHERE MTBC DNA WASN'T DETECTED. 20
FIGURE S10. PLWH HIV-1 DDPCR REPRESENTATIVE QUANTASOFT AMPLIFICATION PLOTS WITH MULTIPLE POSITIVE DROPLETS FOR TOTAL DNA SAMPLES EXTRACTED FROM SPINAL BIOPSY TISSUES. 21
FIGURE S11. RPP30 DDPCR REPRESENTATIVE QUANTASOFT AMPLIFICATION PLOTS FOR TOTAL DNA SAMPLES EXTRACTED FROM SPINAL BIOPSY TISSUES FROM SPINAL TUBERCULOSIS PATIENTS. 22

REFERENCES 23

#### SUPPLEMENTAL METHODS

##### Sex as a biological variable

All individuals consecutively investigated for STB during the study period were informed of the study and given the opportunity to provide consent to participate, irrespective of sex. Data analysis was performed to determine whether there was a sex difference between diagnostic groups.

##### Inclusion and exclusion criteria signs and symptoms of adult patients presenting to our unit for diagnosis.[1]

Back pain with insidious onset, elevated inflammatory markers, TB history/contact, constitutional symptoms, chronic cough, infection with HIV-1, suspicious radiological signs including osteopenia and erosions involving both sides of the affected joint, anterior vertebral height loss, paravertebral or psoas shadow suggesting abscess formation, adjacent endplate changes with preserved disc height, as well as local kyphosis, and were undergoing spinal biopsy (as per clinical indication). Exclusion criteria were age less than 18 years, inability to adhere to study procedures and complete questionnaires, and if there was subsequent isolation of a non-tuberculous *Mycobacterium.*

##### Clinical Data Collection

Clinical data were collected as previously described[1] and were captured in a dedicated REDCap study database. Relevant data included: initial clinical presentation, signs and symptoms, comorbidities including HIV-1 infection, associated peripheral CD4 count and VL, previous TB history, Ziehl-Neelsen stain histology for acid fast bacilli and hematoxylin-eosin stain for histopathology, Xpert MTB/RIF Ultra (GXPU) and associated target gene (*IS1081-IS6110* and *rpoB*) Ct values, MGIT TB culture (TBCUL) time to positivity (TTP), available antibiotic regimen, duration of treatment and outcome, if available, and type of surgical intervention. Medical history of ARV treatment was not readily available.

##### Optimised DNA extraction method from open and CT-guided biopsies

Working in a BSL3 laboratory, containers with biopsy samples in DNA/RNA shield™ were transferred to a Class II biosafety hood and allowed to thaw to room temperature. Tissue samples were cut into four approximately 1 $cm^2$ segments of tissue using a sterile scalpel and each segment placed in a separate sterile ZR BashingBead Lysis Tube (2mm, Zymo Research), comprising of 0.7 ml dry volume 2.0 mm ZR BashingBead lysis matrix (**Supplemental Figure S1-S2**). Due to the smaller nature of the CT-guided biopsy samples, the entire sample was recovered from the DNA/RNA Shield™ (**Supplemental Figure S3**) and placed into one sterile ZR BashingBead Lysis Tube (2mm). 750µl 0.1 x TE buffer was added to each tube and samples were secured in a FastPrep®-24 Classic Bead Beating Grinder and Lysis System (MP Biomedicals™) fitted with a 2ml tube holder assembly and processed at maximum speed for ≥ 5 minutes (1 minute at maximum speed, 5 minutes rest on ice and the cycle was repeated 5 times). Tubes were centrifuged for 1 minute at ≥10,000 g (14,000 g). 500 µl of supernatant from each tube was transferred to a new 1.5 ml Eppendorf tube. In addition, for tissue biopsies, the 4 tubes that contained the 4 segments had the residual 125 µl of supernatant close to the beads transferred and combined into a fifth tube (**Supplemental Figure S1**).

All supernatant samples had DNA extracted using a modified and optimized cetyltrimethylammonium bromide (CTAB) and chloroform-isoamyl alcohol protocol. Briefly, 70 µl of a 10mg/ml lysozyme solution was added and the samples were incubated in a water bath set at 37 °C for 1 hour. The suspension was centrifuged at 20,300 g for 10 minutes to pellet the bacteria and the supernatant was carefully removed. The resulting pellet was re-suspended in 800 µl pre-warmed CTAB/NaCl buffer (4.1g NaCI and 10g CTAB 99+% (ThermoFisher Scientific, Waltham, MA) in 100 ml $H_2O$), containing Proteinase K (10 µl of a 20 mg/ml stock) and incubated at 60 °C with shaking for 1 hour. The tube was gently mixed from time to time by inversion. After 1 hour, 800 µl of chloroform/iso-amyl alcohol (24:1) solution was added in the fume hood and samples were gently mixed by inversion for two minutes. Samples were centrifuged at 14,000 g for 10 minutes at 4°C. The resulting aqueous phase was carefully transferred to a sterile labelled Eppendorf tube, and the rest discarded. 600 µl of ice-cold isopropanol was then added to the tubes, followed by gentle inversion to ensure complete mixing and precipitation at -20°C for 30 minutes. To pellet the DNA, the samples were then centrifuged at 14,000 g for 15 minutes at 4°C after which the supernatant was removed, and the pellet was washed twice with 500 µl ice-cold 70% ethanol by centrifugation at 14,000 g for 15 minutes. The supernatant was carefully removed, and the pellet was air dried. Samples were placed on a heating block at 65°C if residual ethanol was noticed. The pellet was rehydrated overnight in 20µl 0.1 x TE buffer. DNA quantity and purity was assessed using the NanoDrop 2000 Spectrophotometer (ThermoFisher Scientific, Waltham, MA, USA). To ensure equal sensitivity for the detection of *Mtb,* HIV-1 and human genomic DNA (gDNA), all DNA samples were diluted to specified working stock concentrations before addition to ddPCR reactions, enabling uniform DNA amounts added to all assay reactions (**Supplemental Figure S1**).

**Droplet digital PCR for the detection of TB, HIV-1 and Rpp30 in spinal biopsy samples.**

Droplet digital PCR (ddPCR) was performed shortly after successful DNA extraction on 93 samples from spinal biopsy tissues. Duplex and single-plex ddPCR assays were set up using the QX200 AutoDG Droplet Digital PCR system (BioRad Laboratories, Hercules, CA) to detect DNA of two MTBC genetic loci that distinguish MTBC from non-tuberculous mycobacteria; IS6110, a repetitive element that is present in multiple copies throughout the *Mtb* genome thereby aiding in maximizing assay sensitivity, and *rpoB*, a single copy gene known to undergo mutations conferring rifampicin resistance. Notably, a low frequency of *Mtb* clinical strains have been found to lack the IS6110 element and thus reliance on this gene alone for MTBC detection can lead to false negatives.[2] An additional duplex assay was performed to detect DNA of two HIV-1 associated genes, *pol* and *gag*, and a single-plex assay was set-up to detect DNA of *RPP30*, to quantify copies of human gDNA and measure the abundance of human cells within the tissues for data normalization.

For each ddPCR assay, details of oligonucleotides, thermocycling conditions and DNA template amount/reaction are detailed in **Supplemental Tables S5-S8**. A maximum of 1µg DNA was added to MTBC and HIV-1 reactions and 100 ng DNA for *RPP30* reactions. Each assay required a specific volume of template DNA, therefore, prior to ddPCR, each gDNA extract was diluted with nuclease-free water, as follows for the detection of: *MTBC (rpoB and IS6110):* 100ng/µl working stocks; *HIV-1, pol and gag:* 125ng/µl working stocks and *RPP30*: 20ng/µl working stocks. Reactions were prepared by adding 11µl 2X ddPCR Supermix for Probes (no dUTP) (BioRad), 1 µl of prepared Oligomix containing the relevant primers and probes (100 nM each), 0.5µl BanII (NEB), 0.7µl rCutSmart™ Buffer and a specified volume of DNA template.

For MTBC detection: 23.2µl reactions were prepared for analysis containing 10µl (≤1 µg) of DNA template; for HIV-1 detection: 20µl reactions were prepared for analysis containing 8µl (≤ 1ug) of DNA template. For RPP30 detection and quantification, 5µl (~100ng) of DNA template was added, as human cells have two RPP30 gene copies per cell, loading more than 100 ng would cause oversaturation of the droplets. The mastermix was prepared according to **Supplemental Table S7**. ZEN Probes, either FAM or HEX, with a double-quenching system were used to reduce background signal. Positive controls included 0.1 ng H37RV genomic DNA extracted from a 5 ml culture using the extraction method described by Belay *et al* (2021) [3] for the MTBC assay and 1 µl and 5 µl of a 7.5ng/µl stock of genomic DNA isolated from the 8E5 T-cell line obtained from the NIH AIDS reagent program (ARP-95) for the HIV and RPP30 reaction, respectively.

A restriction enzyme digestion was included in the reaction mix using BanII enzyme and rCutSmart™ Buffer (New England Biolabs, R0119L, 1000 U/ml). This is recommended when loading greater than 60 ng gDNA into a reaction to reduce viscosity and improve ddPCR sensitivity. Using GenScript Restriction Enzyme Map Analysis Tools (https://www.genscript.com/tools/restriction-enzyme-map-analysis), it was confirmed that BanII does not cut within either MTBC amplicon. The digest was performed inside the PCR mastermix tube prior to PCR cycling. The reactions were incubated at 37°C for 30 minutes in a Bio-Rad C1000 touch thermocycler to digest the gDNA prior to generating droplets. After gDNA digestion, 40 µL of droplets per well were generated using the autoDG QX200 Droplet Generator according to the manufacturer's instructions. 96-well plates were sealed with foil using the Bio-Rad PX1 PCR plate sealer and PCR was performed using the Bio-Rad C1000 touch thermal cycler. Thermal cycling conditions for each assay are presented in **Supplemental Table S8**.

The QX200 droplet reader was used to analyse the droplets according to manufacturer's instructions. QuantaSoftTM Analysis Pro software version 1.0.596 was employed for data analysis. Positive droplets, containing amplified gene products were discriminated from negative droplets by applying a threshold set at the following amplitudes specific for each assay: *MTBC detection*: amplitude 3000 for channel 1 (detecting the FAM-labelled rpoB probe) and 1500 for channel 2 (detecting the HEX-labelled IS6110 probe), *HIV-1 detection*: amplitude 2500 for channel 1 (detecting the

FAM-labelled pol probe) and 1500 for channel 2 (detecting the HEX-labelled gag probe) and *RPP30 detection*: 1250 for channel 2 (detecting the HEX-labelled RPP30 probe).

Example plots showing thresholds and positive and negative controls for each gene are shown in **Figures S6-S11**. Data from wells with droplet counts <10,000 were excluded from statistical analyses, according to the manufacturer's instructions. The copy number concentration of each sample was reported automatically by the QuantaSoftTM Analysis Pro software. Positive controls and no-template controls (NTC) were included in all assays. Mean droplet count from wells with droplet counts >10,000 and the mean total volume of the droplets measured per reaction, assuming the manufacturer's published volume of 0.85 nL per droplet, are summarized for each assay in Table S5. After quality control (QC) and assessment of ddPCR results, valid data (ie, droplet counts >10,000 per well) were available for 87/93 (94%) of MTBC reactions, 39/41 (95%) of HIV-1 reactions and 93 (100%) of *RPP30* reactions. For MTBC and HIV-1 samples where less than 1 µg of total DNA was added to the reaction, the resulting copies/20 µl generated by the software had a normalization factor applied, so that results represented 1 µg of input DNA to match all other samples.

Data measured as pathogen DNA copies/µl, were normalized to *RPP30* DNA copies (present at two copies/nucleated cell x 10 to normalise 100 ng input DNA for *RRP30* reaction vs. 1 µg input for pathogen gene reactions) to represent pathogen gene copies/million human nucleated cells, using the equation below, thus allowing pathogen abundance comparison across varying sample tissue types (20).

*Gene copies/million cells = (pathogen gene copies/µl)/((RPP30 copies/µl)/2 x 10) x 1 million*

**Accuracy and calculation of limit of detection of MTBC DNA ddPCR assay**

To determine the lowest concentration of *Mtb* DNA that can be reliably and consistently measured by the ddPCR assay, DNA extracted from the laboratory strain *Mtb* H37Rv was subjected to a 10-fold dilution series and technical replicates from 10 pg to 10 fg per 20µl reaction volume were assayed. Genomic DNA (gDNA) was extracted as described above from a concentrated and heat-killed (two hours in 80ºC water bath) pellet from a 5ml culture of laboratory strain *Mtb* H37Rv grown in 7H9/OADC broth (BD Middlebrook). The resulting DNA was reconstituted in nuclease-free water and stored at -20°C. Serial 10-fold dilutions of the DNA stock ranging from 10pg (~2000 *Mtb* DNA copies/reaction) down to 10fg (~2 Mtb DNA copies /reaction) were assayed with the duplex ddPCR assay for the detection of both *rpoB* and *IS6110* (**Supplementary Figure S5 & S6**). Resulting data were utilized for calculating the limit of blank (LOB) and the limit of detection (LOD), considering the blank no template controls (NTCs), where no DNA template was added, as previous [3]. The LOD is the lowest sample analyte concentration likely to be reliably distinguished from the LOB and at which detection is feasible. The limit of detection (LOD) for each MTBC gene was calculated using the following formula:

*Limit of detection = limit of blank + 1.645 x SD(low concentration sample) where:*
*Limit of blank= mean (blank) + 1.645 x SD (blank)*

This guided the appropriate statistical analysis of the detection of both *rpoB* and *IS6110* in spinal biopsy DNA samples. Both *rpoB* and *IS6110* were readily detectable using 10 fg MTBC DNA with LODs determined as 2.29 copies/20µl reaction for *rpoB* and 9.67 copies/20µl reaction for multi-copy *IS6110*.

Determination of the limit of blank and limit of detection were not standard practice for the detection of HIV targets. Rather, the maximum concentration of the Poisson confidence intervals for the negative and non-template control samples was used as the concentration at or below which signal from an experimental sample was considered below background.

#### SUPPLEMENTAL TABLES

**Table S1. Histopathological or other diagnosis for the nine patients not diagnosed with STB.**

| Patient Study ID | HIV Status | Previous PTB | Histopathological diagnosis | Outcome | MTBC ddPCR |
|---|---|---|---|---|---|
| MUTI010 | Positive | No | Fibrocollagenous tissue with chronic inflammation. | | Negative |
| MUTI013 | Negative | Yes | Non-necrotizing granulomatous inflammation | | Negative |
| MUTI015 | Negative | No | Chronic osteomyelitis | | Negative |
| MUTI017 | Negative | Yes | Unknown pathology, negative for malignancy* | | *rpoB+IS6110+* |
| MUTI020 | Negative | Yes | Mild acute inflammation | Deceased | *rpoB+IS6110+* |
| MUTI021 | Negative | No | Abscess | | Negative |
| MUTI022 | Negative | No | Unknown pathology* | | Negative |
| MUTI027 | Positive | Yes | Consistent with fracture site, negative for malignancy. | Deceased | *rpoB+IS6110-* |
| MUTI029 | Positive | Yes | No histology report | | *rpoB+IS6110-* |

*No pathologic diagnosis.

**Table S2. Per patient detection of *Mtb*, HIV-1 and human DNA copies in spinal tissues, stratified by STB definition and HIV status.**

| Patient MUTI study ID | STB definition | HIV-1 status | HIV-1 Viral Load (copies/ml) | Previous TB history | Biopsy method | Tissue description | Total DNA extracted (µg) | Total human cells ($x10^6$) | Median (IQR) total human cells/open biopsy segment ($x10^6$) | Number of MTBC positive tissue segments | Total MTBC *rpoB* copies/ biopsy | Total MTBC *IS6110* copies/ biopsy | Number of HIV-1 positive tissue segments | Total HIV-1 *Pol* copies/ biopsy | Total HIV-1 *Gag* copies/ biopsy |
|---|---|---|---|---|---|---|---|---|---|---|---|---|---|---|---|
| **13** | Not STB | Negative | | Yes | CT-guided Needle | 2-7x1mm | 2.2 | 0.048 | - | 0/1 (0%) | 0 | 0 | - | - | - |
| **15** | Not STB | Negative | | No | Open surgery | Soft bone | 63.7 | 4.66 | 1.22 (0.28 - 8.37) | 0/4 (0%) | 0 | 0 | - | - | - |
| **17** | Not STB | Negative | | Yes | Open surgery | Soft bone | 156.7 | 9.94 | 1.91 (0.10 - 1.26) | 1/5 (20%) | 85.9 | 386.6 | - | - | - |
| **20** | Not STB | Negative | | Yes | Open surgery | Soft bone | 25.6 | 3.35 | 2.36 (1.76 - 1.27) | 2/5 (40%) | 122.1 | 254.0 | - | - | - |
| **21** | Not STB | Negative | | No | Open surgery | Soft bone | 227.8 | 30.4 | 4.66 (3.21-10.40) | 0/5 (0%) | 0 | 0 | - | - | - |
| **22** | Not STB | Negative | | No | Open surgery | Hard bone | 32.1 | 0.89 | 1.84 (1.32 -1.91) | 0/4 (0%) | 0 | 0 | - | - | - |
| **10** | Not STB | Positive | LDL | No | CT-guided Needle* | 2-7x1mm | 0.7 | 0.06 | - | 0/1 (0%) | 0 | 0 | 1/1 (100%) | 0 | 16.4 |
| **27** | Not STB | Positive | 676 | Yes | Open surgery | Hard bone | 96.8 | 9.88 | 2.12 (0.71 - 2.74) | 1/5 (20%) | 44.4 | 0 | 5/5 (100%) | 926.5 | 1951.5 |
| **29** | Not STB | Positive | 60 | Yes | Open surgery | Hard bone | 19.3 | 3.17 | 0.44 (0.11 - 0.71) | 1/4 (25%) | 59.0 | 0 | 4/5 (80%) | 61.5 | 982.2 |
| **11** | Xpert Ultra confirmed | Negative | | Yes | Open surgery | Soft bone | 156.7 | 24.7 | 3.73 (2.80- 8.37) | 5/5 (100%) | 2962.0 | 16497.8 | - | - | - |
| **23** | Xpert Ultra confirmed | Negative | | No | Open surgery* | Other | 2.8 | 0.32 | 0.036 (0.03- 0.06) | 3/5 (60%) | 33.1 | 30.2 | - | - | - |
| **14** | Xpert Ultra confirmed | Positive | NA | No | CT-guided Needle* | 2-7x1mm | 2.5 | 0.14 | - | 1/1 (100%) | 24.4 | 579.0 | 1/1 (100%) | 162.7 | 291.6 |
| **16** | Xpert Ultra confirmed | Positive | LDL | Yes | CT-guided Needle* | 2-7x1mm | 8.0 | 2.31 | - | 0/1 (0%) | 0 | 0 | 1/1 (100%) | 209.9 | 386.2 |
| **18** | Xpert Ultra confirmed | Positive | 49959 | Yes | CT-guided Needle* | 2-7x1mm | 2.4 | 2.35 | - | 1/1 (100%) | 51.6 | 218.4 | 1/1 (100%) | 187.2 | 0.0 |
| **19** | Xpert Ultra confirmed | Positive | 1137 | Yes | CT-guided Needle* | 2-7x1mm | 0.7 | 0.04 | - | 1/1 (100%) | 7.9 | 146.7 | 1/1 (100%) | 25.9 | 20.7 |
| **9** | Culture confirmed | Negative | | No | CT-guided Needle | 2-7x1mm | 1.5 | 0.11 | - | 1/1 (100%) | 41.4 | 49.8 | - | - | - |
| **24** | Culture confirmed | Negative | | No | Open surgery | Other | 233.6 | 34.1 | 6.42 (5.52 - 8.67) | 5/5 (100%) | 1259.7 | 1192.4 | - | - | - |
| **25** | Culture confirmed | Negative | | No | Open surgery | Hard bone | 64.3 | 7.46 | 0.84 (0.27 -2.44) | 3/4 (75%) | 292.2 | 4617.1 | - | - | - |
| **30** | Culture confirmed | Negative | | No | Open surgery | Soft bone | 44.3 | 6.99 | 0.62 (0.46 - 1.36) | 1/4 (25%) | 30.8 | 0 | - | - | - |
| **28** | Culture confirmed | Positive | 446 | Yes | CT-guided Needle* | 2-7x1mm | 0.7 | 0.02 | - | 1/1 (100%) | 49.2 | 184.5 | 1/1 (100%) | 5.3 | 105.6 |
| **2** | Culture confirmed | Positive | 372 | Yes | Open surgery* | Soft bone | 155.2 | 26.3 | 5.12 (3.09 - 8.13) | 5/5 (100%) | 34272.9 | 39907.3 | 5/5 (100%) | 1218.7 | 7537.6 |
| **26** | Culture confirmed | Positive | 50 | Yes | Open surgery | Hard bone | 149.5 | 10.1 | 1.98 (1.84 - 2.60) | 4/5 (80%) | 5434.9 | 31420.4 | 4/4(100%) | 1555.0 | 2350.4 |
| **3** | Culture confirmed | Positive | LDL | Yes | Open surgery* | Soft bone | 241.6 | 16 | 0.57 (0.22- 1.79) | 1/4 (25%) | 51.3 | 58.4 | 3/4 (75%) | 15.2 | 579.8 |
| **31** | Culture confirmed | Positive | LDL | Yes | Open surgery | Soft bone | 306.5 | 36.9 | 7.91 (5.39 - 9.10) | 5/5 (100%) | 59144.7 | 1001775.2 | 2/3 (67%) | 107.4 | 372.4 |
| **7** | Culture confirmed | Positive | NA | No | Open surgery | Soft bone | 207.4 | 13 | 1.84 (0.81 - 4.13) | 3/5 (60%) | 1427.6 | 5985.5 | 5/5 (100%) | 7789.8 | 8298.9 |

Total DNA extracted was calculated by adding total amounts of DNA extracted from each biopsy segment for open surgeries (n=5; n=3 or 4 indicates the samples which had one or two segments excluded due to invalid results with <10,000 droplets detected) or the total extracted from the core percutaneous CT-guided needle biopsy (n=1). Total number of cells was calculated by considering *RPP30* copies/reaction, two *RPP30* copies/nucleated cell and total DNA extracted. Number of MTBC or HIV-1 DNA positive tissue segments indicates the proportion of segments with either pathogen-specific gene detected for that patient. Total gene copies per biopsy tissue were calculated by considering gene copies/reaction/segment and total DNA extracted/segment, and total segments were summed per patient. LDL: Lower than detectable limit/undetectable; NA: not applicable. *: On TB treatment at the time of biopsy (n=9).

**Table S3. Mean droplet count and mean total volume of droplets for each ddPCR assay**

| ddPCR assay | Mean droplet count/reaction (SD, range) | Mean total volume of droplets/reaction (SD, range) |
|---|---|---|
| **MTBC** ***rpoB/IS6110*** **assay** | 17,186 (SD: 1455, range: 12,593 to 21,402) | 14.3 µl (SD: 1.3 µl, range: 8.58 to 15.96 µl) |
| **HIV-1** ***pol/gag*** **assay** | 16,775 (SD: 1459, range: 10,799 to 18,842) | 14.3 µl (SD: 1.2 µl, range: 9.17 to 16.01 µl) |
| **Human** ***RPP30*** **assay** | 17,604 (SD: 1105, range: 10,318 to 19,020) | 15.0 µl (SD: 0.9 µl, range: 8.77 to 16.16 µl) |

**Table S4. Detection of MTBC DNA by patient HIV-1 status.**

| | | **MTBC DNA (*rpoB* or *IS6110*)** | | |
|---|---|---|---|---|
| | | **Detected** | **Undetected** | **p-value** |
| **HIV-1 infection status** | **HIV-1-uninfected** | 8/25 (32%) | 4/25 (16%) | 0.3783 |
| | **HIV-1-infected** | 11/25 (44%) | 2/25 (8%) | |

*P*-values were calculated by Fisher's exact test.

**Table S5. Oligonucleotide information, MTBC DNA ddPCR assays.**

| Gene target | Oligonucleotide name | Oligonucleotide sequence | Reporter, Quencher | oligos nM | Amplicon length, bp | Reference |
|---|---|---|---|---|---|---|
| *IS6110*[A] | *Forward primer, IS6110* | AGAAGGCGTACTCGACCTGA | - | 100 | 157 | 3 |
| | *Reverse primer, IS6110* | GATCGTCTCGGCTAGTGCAT | - | 100 | | |
| | *HEX-labelled probe, IS6110* | AGGCAGGCATCCAACCG | HEX, BHQ1 (3IABkFQ) | 100 | | |
| *rpoB*[B] | *Forward primer, rpoB* | CAAAACAGCCGCTAGTCCTAGTC | - | 100 | 84 | |
| | *Reverse primer, rpoB* | AAGGAGACCCGGTTTGGC | - | 100 | | |
| | *FAM-labelled probe, rpoB* | AGTCGCCCGCAAAGTTCCTCGAA | FAM,BHQ1 (3IABkFQ) | 100 | | |

[A]Base pair locations of the IS6110 motifs on the bacterial genome are 889985...890141, 1542342...1542186, 1988093...1987937, 1997065...1997221, 2366378...2366534, 2430507...2430351, 2550978...2551134, 2636541...2636697, 2785005...2784849, 2973073...2973229, 3120914...3120758, 3552194...3552350, 3553677...3553833, 3711346...3711502, 3795448...3795292, 3891743...3891899. Sequence information based on *Mycobacterium tuberculosis* H37Rv, NC_018143. [B]Base pair location of the rpoB motif on the bacterial genome is 759827… 759910. Sequence information based on *Mycobacterium tuberculosis* H37Rv, NC_018143.

**Table S6. Oligonucleotide information for HIV-1 and RPP30 ddPCR assays.**

| Gene target | Oligonucleotide name | Oligonucleotide sequence | Reporter, Quencher | HXB2 nucleotide position (relative to start of gene region) | Reference |
|---|---|---|---|---|---|
| *pol* | *pol Fwd 6* | TCG GGT TTA TTA CAG AGA CAG CAG AGA | - | 2814 - 2840 | Modified from [4] |
| | *pol Rev 4* | AGC ICC TGC CAT CTG TTT TCC AT | - | 2957 - 2979 | |
| | *pol probe 3 FAM ZEN* | /56-FAM/AAG GAC CAG/ZEN/CCA ARC TAC TCT GGA AAG GTG/3IABkFQ/ | FAM, BHQ1 (3IABkFQ) | 2852 - 2881 | |
| *gag* | *gag Fwd* | GTT GGA GGA CAT CAA GCA GCC ATG CA | - | 571 - 596 | Modified from [4] |
| | *gag Rev* | TTC CTG CTA TGT CAC TTC CCC T | - | 694 - 715 | |
| | *gag probe HEX ZEN* | /5HEX/ACC ATC AAT/ZEN/GAR gag GCT GCA GAA TGG GA/3IABkFQ/ | HEX, BHQ1 (3IABkFQ) | 610 - 638 | |
| *RPP30* | *RPP30 Fwd* | GAT TTG GAC CTG CGA GCG | - | - | NA |
| | *RPP30 Rev* | GCG GCT GTC TCC ACA AGT | - | - | |
| | *RPP30 probe HEX ZEN* | /5HEX/CTG ACC TGA/ZEN/AGG CTC T/3IABkFQ/ | HEX, BHQ1 (3IABkFQ) | - | |

#### Table S7. Reaction setup per ddPCR assay (MTBC, HIV-1 and RPP30 detection)

| ***rpoB /IS6110*** | **Stock Concentration** | **Final concentration** | **Vol (µl)** |
|---|---|---|---|
| Supermix | 2x | 1x | 11 |
| Oligomix | 22x | 1x | 1 |
| BanII | 10U/µl | 0.25U/µl | 0.5 |
| CutSmart buffer | 10x | 0.35x | 0.7 |
| dH20 | - | - | 0 |
| Template DNA | - | - | 10 |
| | | ***Total*** | ***23.2*** |
| ***Gag/Pol*** | **Stock Concentration** | **Final concentration** | **Vol (µl)** |
| Supermix | 2x | 1x | 10 |
| FAM Forward Primer | 50µM | 0.45µM | 0.18 |
| FAM Reverse Primer | 50µM | 0.45µM | 0.18 |
| FAM Probe | 25µM | 0.25µM | 0.2 |
| HEX Forward Primer | 50µM | 0.45µM | 0.18 |
| HEX Reverse Primer | 50µM | 0.45µM | 0.18 |
| HEX Probe | 25µM | 0.25µM | 0.2 |
| BanII | 10U/µl | 0.25U/µl | 0.5 |
| CutSmart buffer | 10x | 0.35x | 0.7 |
| dH20 | - | - | 0 |
| Template DNA | - | - | 8 |
| | | ***Total*** | ***20.32*** |
| ***RPP30*** | **Stock Concentration** | **Final Concentration** | **Vol (µl)/rxn** |
| Supermix | 2x | 1x | 10 |
| HEX Forward Primer | 10µM | 0.45µM | 0.9 |
| HEX Reverse Primer | 10µM | 0.45µM | 0.9 |
| HEX Probe (ZEN) | 10µM | 0.25µM | 0.5 |
| BanII | 10U/µl | 0.25U/µl | 0.5 |
| CutSmart buffer | 10x | 0.35x | 0.7 |
| dH20 | - | - | 1.5 |
| Template DNA | - | - | 5 |
| | | ***Total*** | ***20*** |

#### Table 8. Thermal cycling conditions, ddPCR assays

| Assay name | Mastermix (Manufacturer) | Enzyme activation | Denaturation | Primer annealing and extension | Droplet stabilisation | Maintenance | Ramp rate |
|---|---|---|---|---|---|---|---|
| *rpoB, IS6110* | ddPCR: 2 x dPCR SuperMix for probes (No dUTP) (BioRad) | 95ºC 10 min | 95ºC 30 secs | 55ºC 1 min | 95ºC 10 min | 4ºC | 2.5ºC/s |
| | | | **39 cycles** | | | | |
| *Pol/gag* and *Rpp30* | | 95ºC 10 min | 94ºC 30 secs | 60ºC[A] 1 min | 98ºC 10 min | 4ºC | 2.5ºC/s |
| | | | **50 cycles** | | | | |

[A]59ºC is best for *gag* primers, 60ºC is best for *pol* primers. *RPP30* works well at both temperatures.

#### SUPPLEMENTAL FIGURES

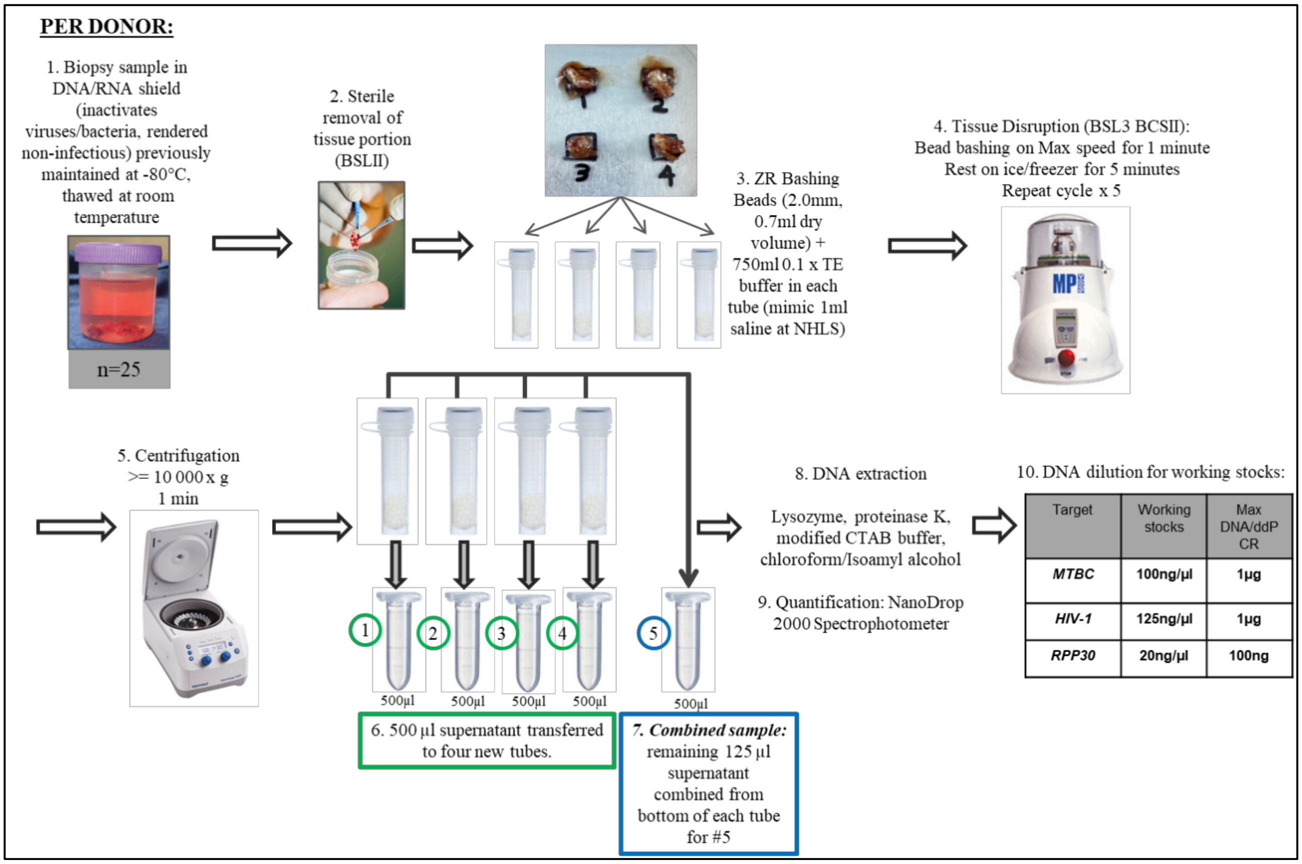

**Figure S1. Total DNA extraction from spinal biopsy specimens**. Open biopsy tissues were cut into four segments for DNA extraction. CT-guided biopsy specimens (not shown here) were loaded into a single tube for extraction, compared to four different tubes for open biopsies.

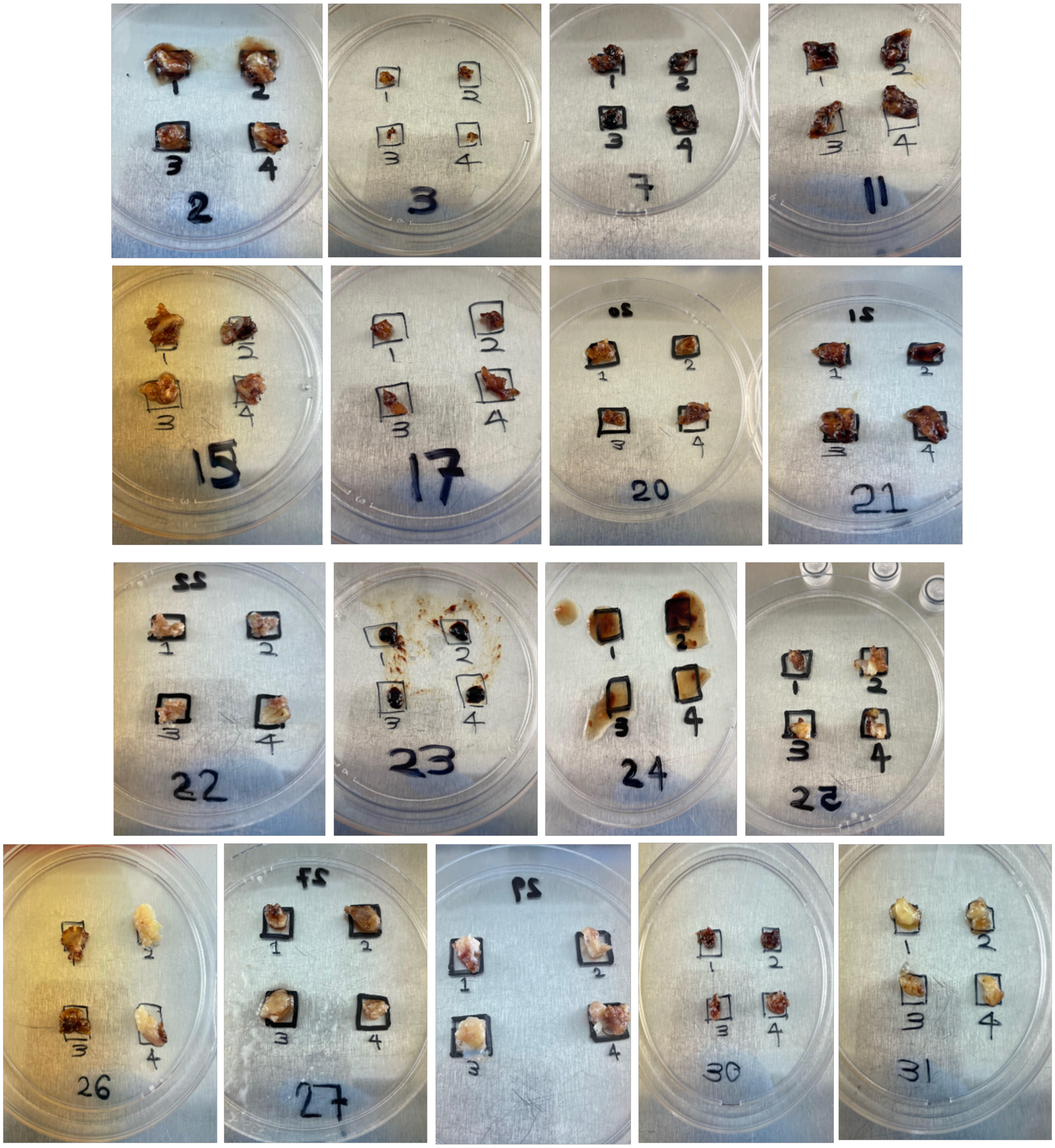

**Figure S2. Representative images of segments of open spinal biopsies with diseased hard bone (white), soft bone (maroon) and other diseased tissue (MUTI024 and MUTI023) (n=17)**.

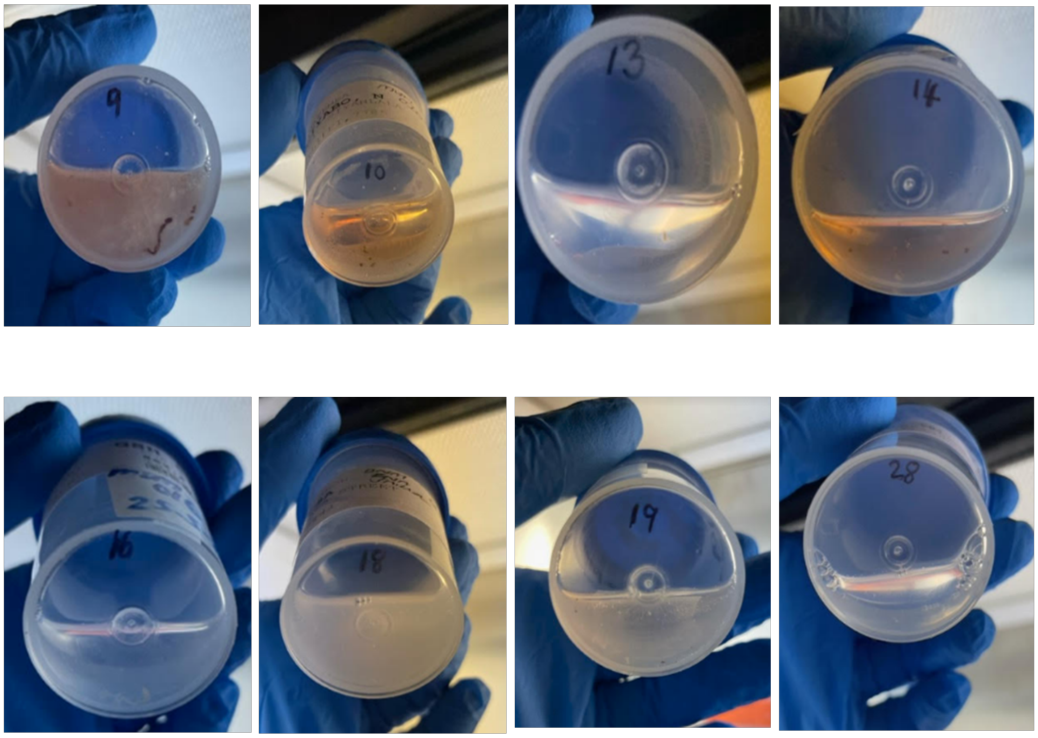

**Figure S3. Representative images of percutaneous CT-guided core biopsies (n=8)**.

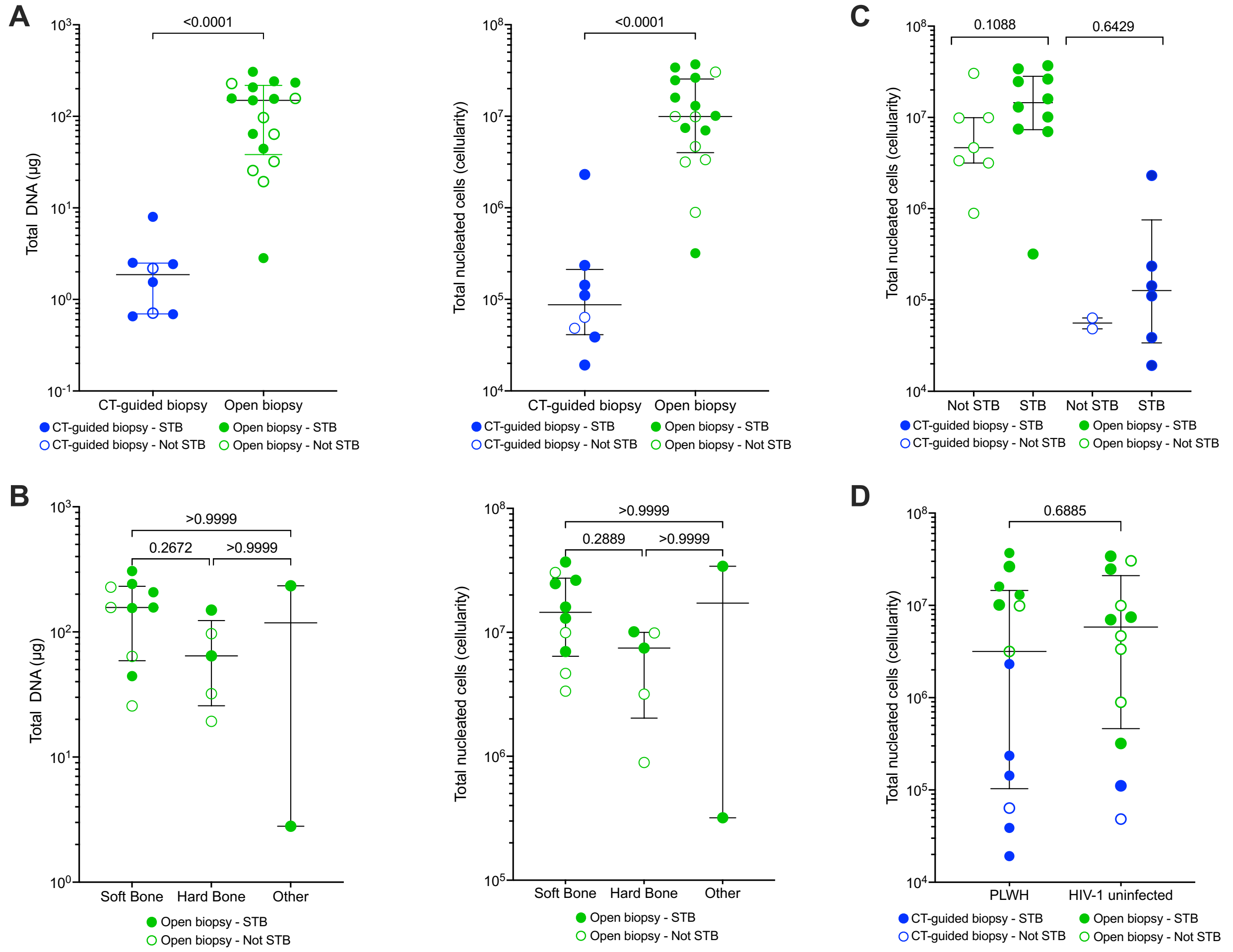

**Figure S4. Total DNA extracted and cellularity of each patients' spinal biopsy tissue, stratified by biopsy method, tissue type, spinal TB diagnosis and HIV status. A**) Total DNA (µg) extracted and total nucleated cells (cellularity) by biopsy method. **B**) Total DNA (µg) extracted and total nucleated cells (cellularity) by open biopsy tissue type. (**C**) Total nucleated cells (cellularity), coloured by biopsy method and stratified by spinal TB (STB: culture confirmed or Xpert Ultra confirmed) diagnosis or **D**) HIV-1 status. Y axes were log transformed. n=8 CT-guided patients and n=17 open biopsy patients. Each dot represents a patient. Total nucleated cells (cellularity), was calculated based on two copies of *RPP30*/nucleated human cell. Open biopsy results represent the sum of the five segments. Median (IQR), two group analysis: Mann-Whitney test, three group analysis: Kruskal-Wallis Test with Dunn's multiple comparison test. CT: Computed tomography; PLWH: People living with HIV-1.

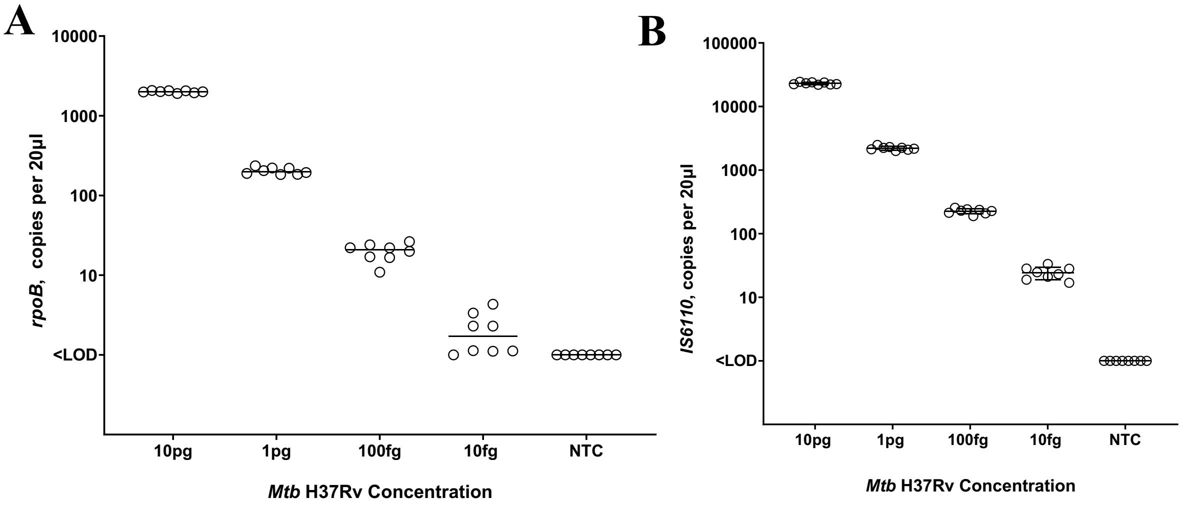

**Figure S5. MTBC gene copy number per 20µl well for a dilution series of H37Rv Mtb DNA.** A) *rpoB*, B) *IS6110.* H37Rv *Mtb*-derived DNA was assayed ranging from 10pg (~2000 *Mtb* DNA copies/reaction) down to 10fg (~2 *Mtb* DNA copies /reaction).

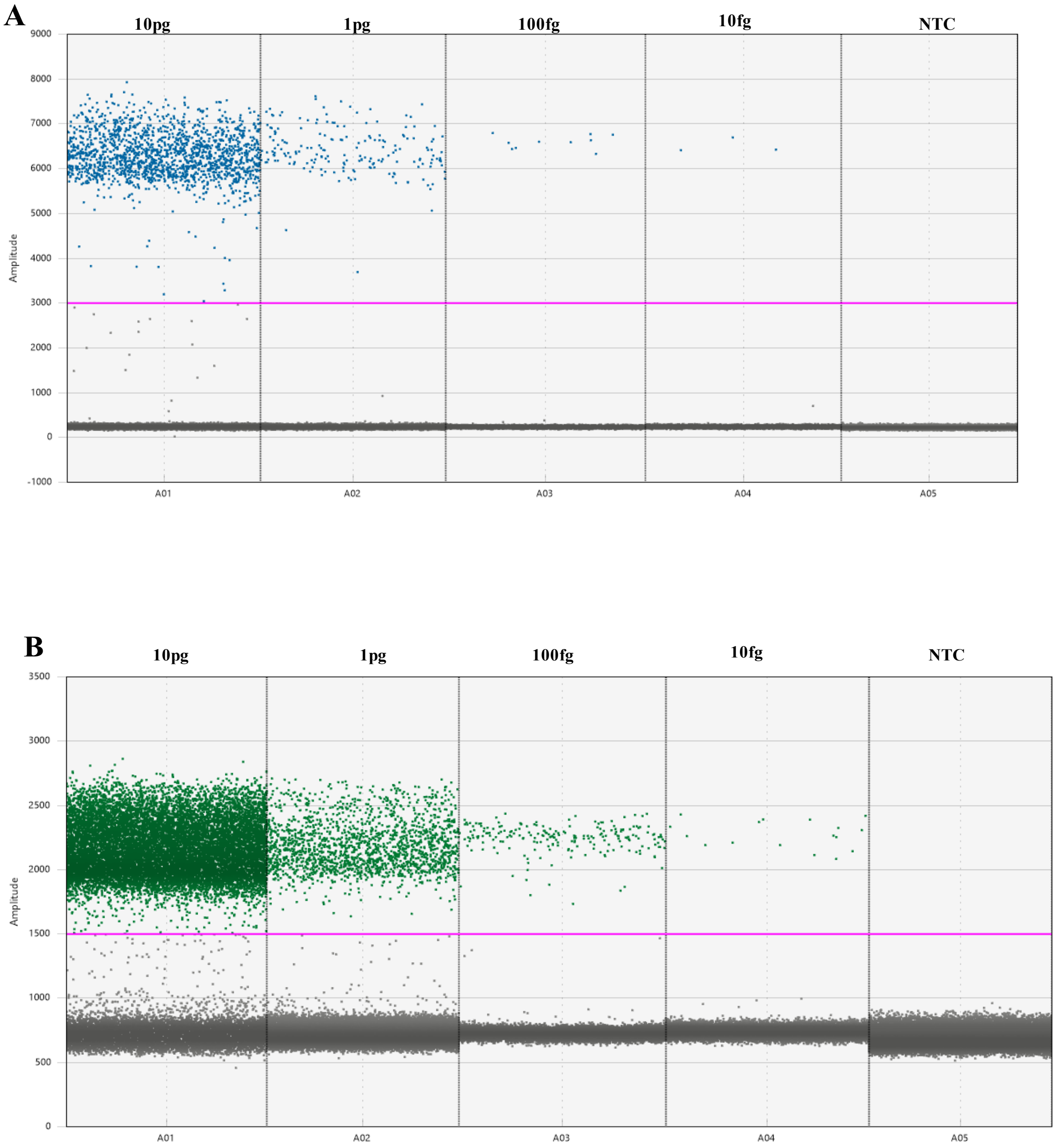

**Figure S6. Representative Quantasoft ddPCR amplification plots from Mtb H37Rv DNA dilution series.** A) *rpoB* (FAM-labelled probe, channel 1, threshold 3000) B) and *IS6110* (HEX-labelled probe, channel 1, threshold 1500). 10pg (~2000 Mtb DNA copies/reaction) 10-fold diluted to 10fg (~2 Mtb DNA copies /reaction). NTC, no template control.

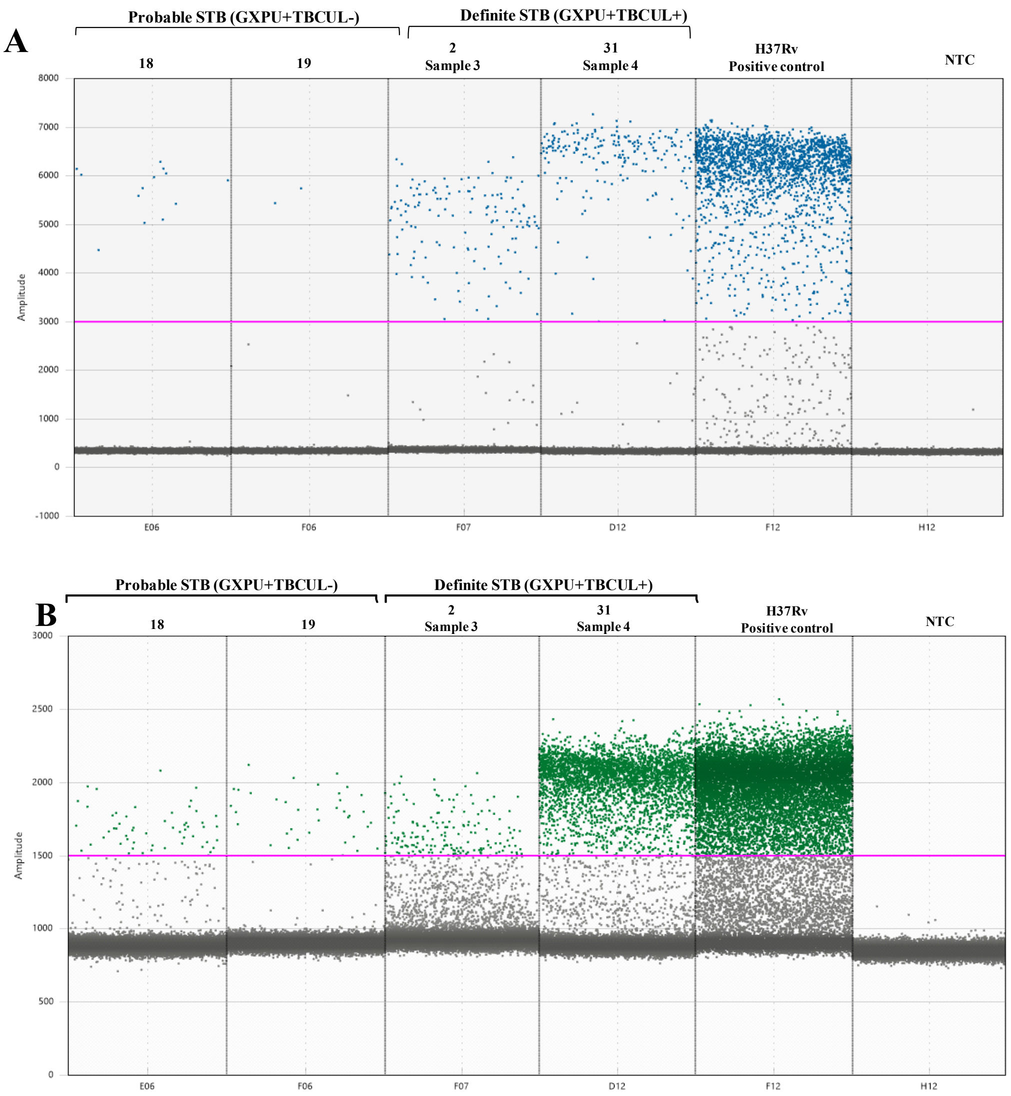

**Figure S7. Probable and Definite STB spinal tuberculosis patients MTBC ddPCR representative Quantasoft amplification plots with multiple positive droplets for total DNA samples extracted from spinal biopsy tissues**. A) *rpoB* (FAM-labelled probe, channel 1, threshold 3000), and B) *IS6110* (HEX-labelled probe, channel 1, threshold 1500). Patient number indicated above column. *NTC, no template control. STB; Spinal Tuberculosis, GXPU: Xpert MTB/RIF Ultra, HIST: Histology.* TB diagnostic definitions: *STB* (TBCUL+GXPU+); *Probable STB* (TBCUL-GXPU+HIST+) ; *NOT STB* (TBCUL-GXPU-HIST-).

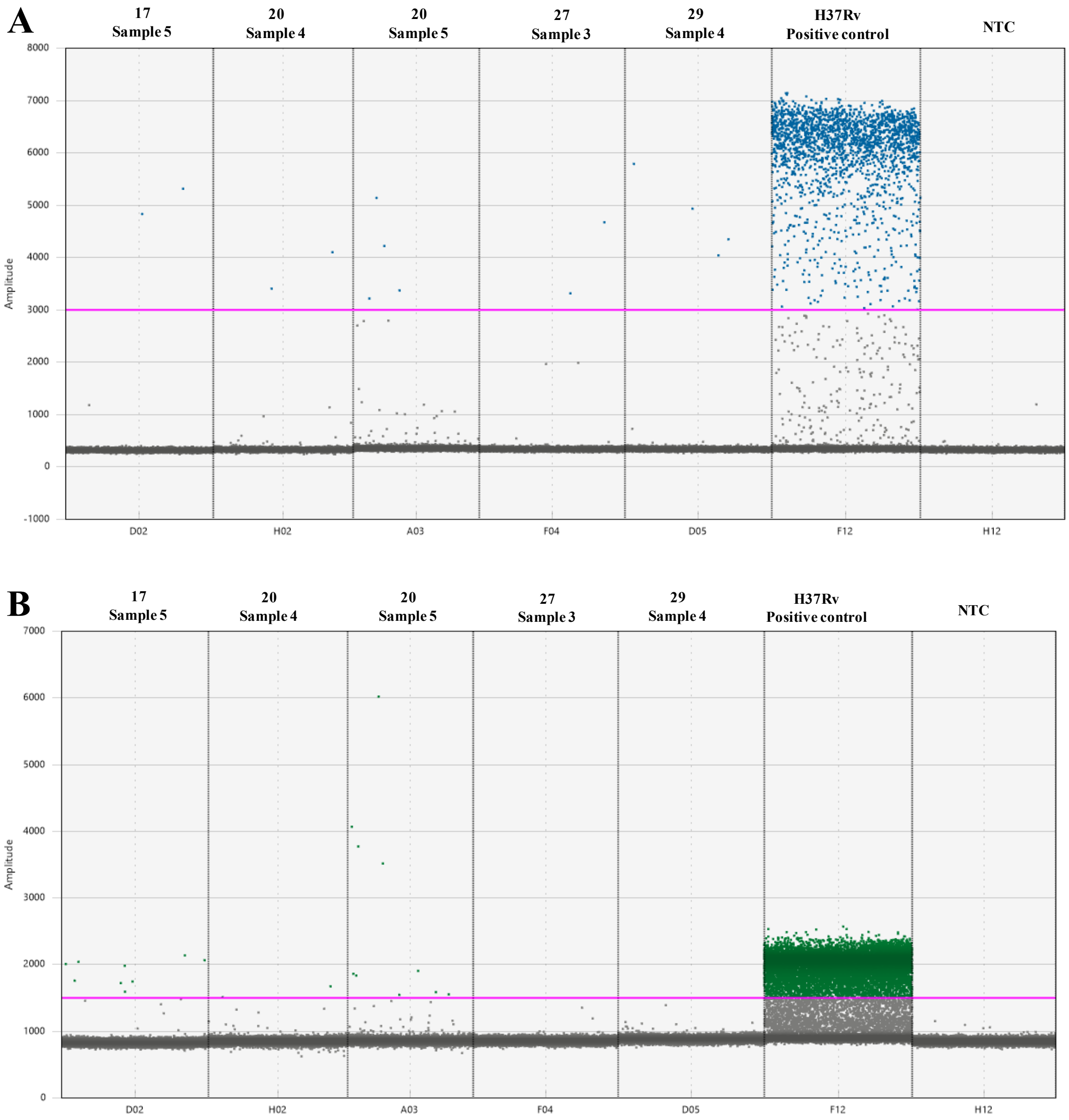

**Figure S8. Not STB (TBCUL-GXPU-HIST-) patients (with previous TB history) MTBC ddPCR representative Quantasoft amplification plots with multiple positive droplets for total DNA samples extracted from spinal biopsy tissues.** A) *rpoB* (FAM-labelled probe, channel 1, threshold 3000) B) and *IS6110* (HEX-labelled probe, channel 1, threshold 1500), for n=5 biopsy segments from n=4 Not STB patients (#17, #20, #27, #29). Patient number indicated above column. *NTC, no template control.*

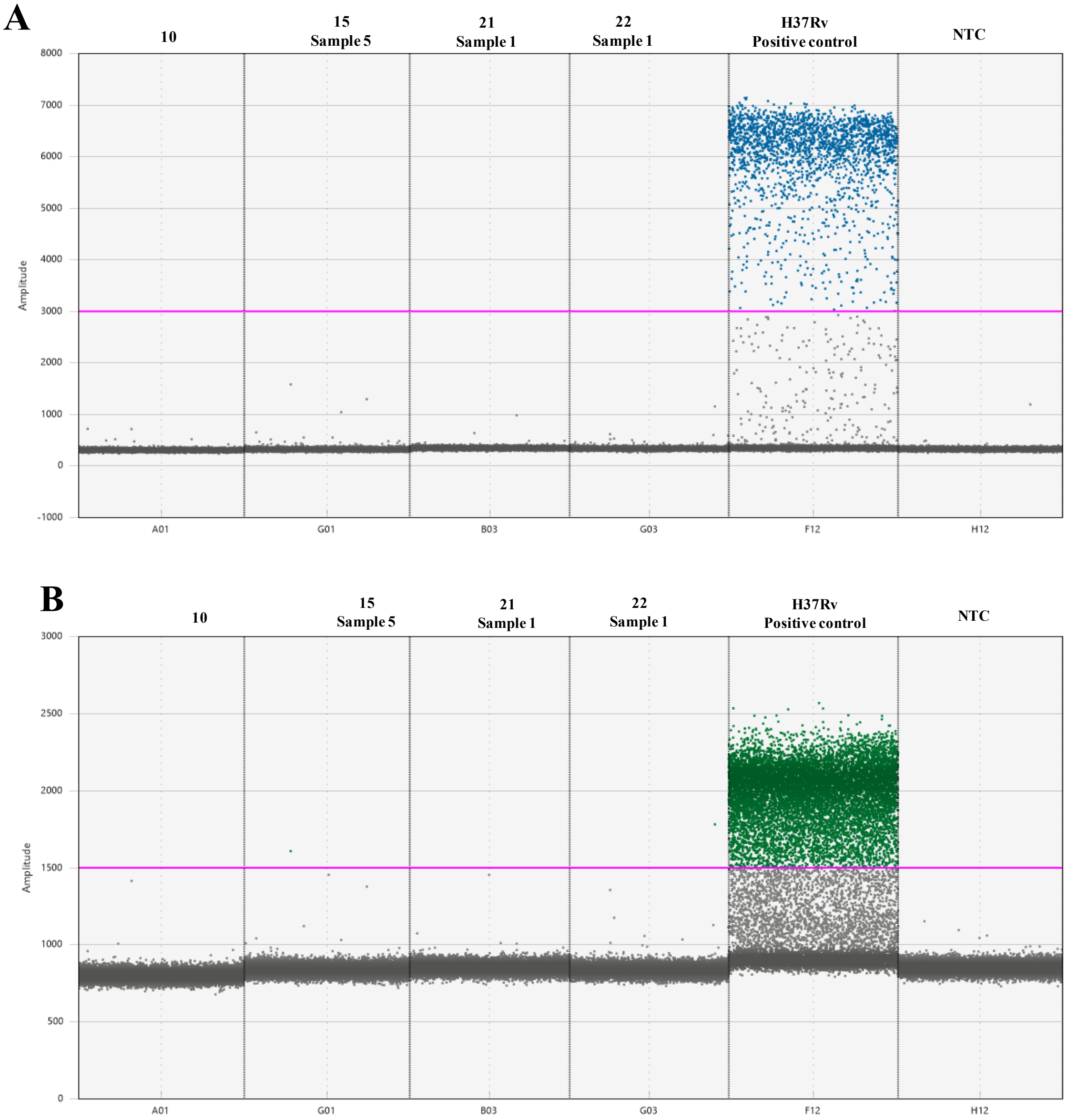

**Figure S9. Not STB (TBCUL-GXPU-HIST-) patients (no previous TB history) MTBC ddPCR representative Quantasoft amplification plots for total DNA samples extracted from spinal biopsy tissues where MTBC DNA wasn't detected.** A) *rpoB (*FAM-labelled probe, channel 1, threshold 3000) B) and *IS6110* (HEX-labelled probe, channel 1, threshold 1500). Two positive dots (15, Sample 5 and 22 Sample 1) not called with confidence by the software. *NTC, no template control.*

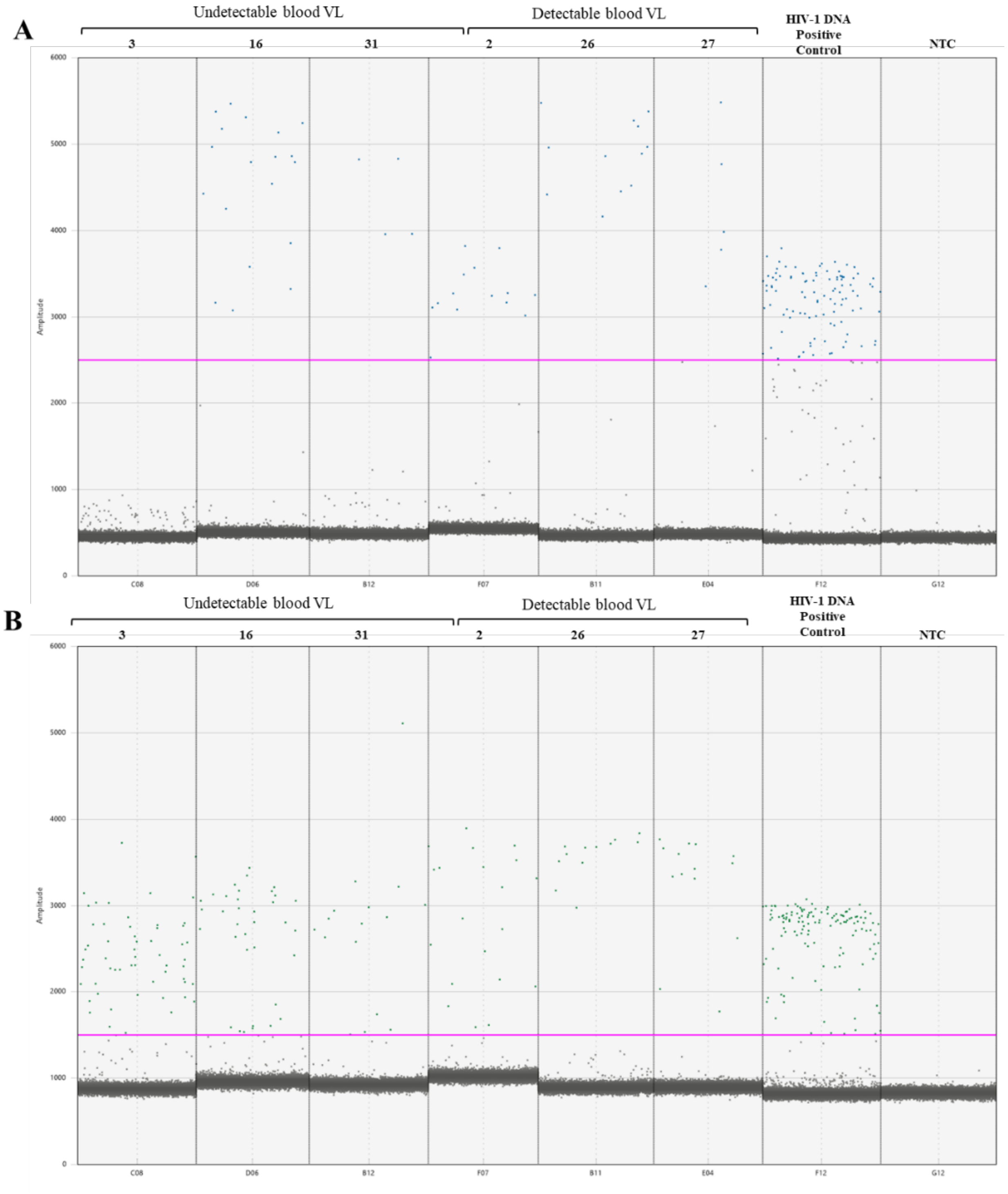

**Figure S10. PLWH HIV-1 ddPCR representative Quantasoft amplification plots with multiple positive droplets for total DNA samples extracted from spinal biopsy tissues.** A) pol FAM-labelled probe, channel 1, threshold 2500) B) and gag (HEX-labelled probe, channel 2, threshold 1500). NTC, no template control.

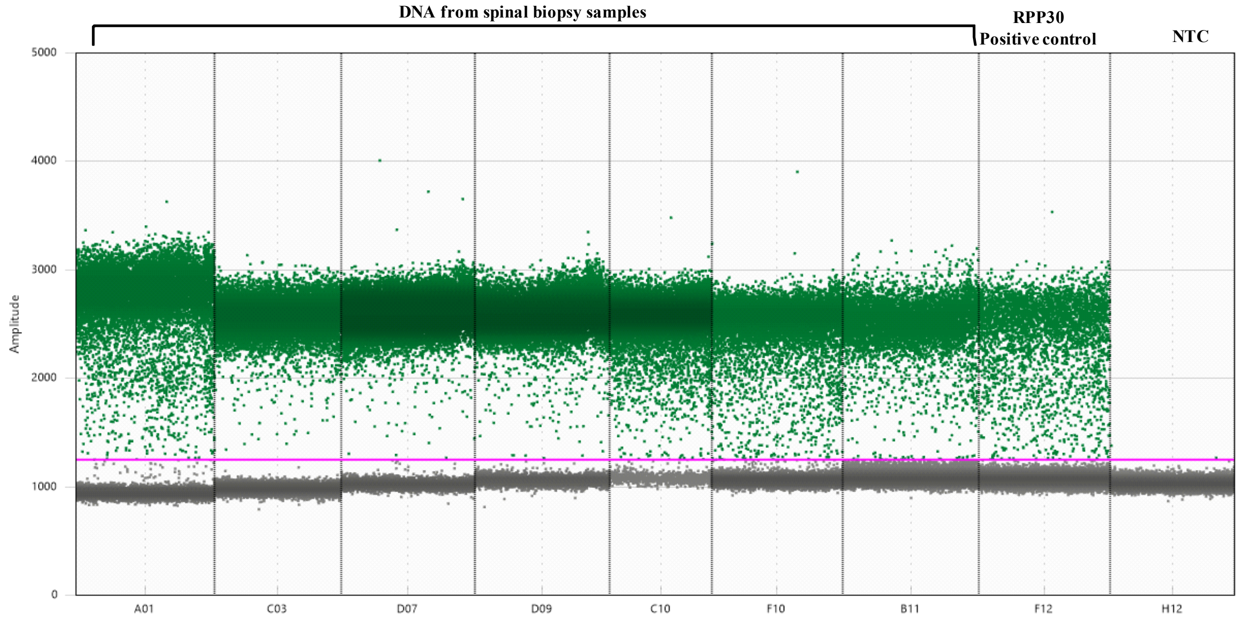

**Figure S11. RPP30 ddPCR representative Quantasoft amplification plots for total DNA samples extracted from spinal biopsy tissues from spinal tuberculosis patients.** *RPP30* (HEX-labelled probe, channel 2, threshold 1250). *NTC, no template control.*
